# FSD1: a plastidial, nuclear and cytoplasmic enzyme relocalizing to the plasma membrane under salinity

**DOI:** 10.1101/2020.03.24.005363

**Authors:** Petr Dvořák, Yuliya Krasylenko, Miroslav Ovečka, Jasim Basheer, Veronika Zapletalová, Jozef Šamaj, Tomáš Takáč

**Author notes:** Correspondence: Tomáš Takáč.

## Abstract

Here, we aimed to resolve the developmental expression and subcellular localization of *Arabidopsis* iron superoxide dismutase FSD1, which belongs to the family of superoxide dismutases (SODs), prominent enzymes decomposing superoxide anion and determining abiotic stress tolerance. We found that *fsd1* knockout mutants exhibit reduced lateral root number and that this phenotype was complemented by *proFSD1::GFP:FSD1* and *proFSD1::FSD1:GFP* constructs. Light sheet fluorescence microscopy revealed a temporary accumulation of FSD1-GFP at the site of endosperm rupture during seed germination. In emerged roots, FSD1-GFP showed the highest abundance in cells of the lateral root cap, columella, and endodermis/cortex initials. The largest subcellular pool of FSD1-GFP was localized in the plastid stroma, while it was also located in the nuclei and cytoplasm. FSD1 is crucial for seed germination and salt stress tolerance, which is tightly coupled with FSD1-GFP subcellular relocation to the plasma membrane. FSD1 is most likely involved in superoxide decomposition in the periplasm. This study suggests a new osmoprotective function of SODs in plants.

---

Plants, as aerobic organisms, have to deal with the harmful by-products of oxidative metabolism named reactive oxygen species (ROS), physiologically produced in organelles (chloroplasts, mitochondria, peroxisomes, glyoxysomes), cytosol, and apoplast. Moreover, ROS play regulatory and signalling roles during plant development and response to environmental challenges^1-5^. To regulate ROS levels, plants have developed adaptations and scavenging machineries^6,7^. Due to compartmentalized ROS production, the antioxidant system is present in different cellular compartments. However, the importance of developmental regulations, tissue-specific expression patterns, and subcellular localizations of antioxidant compounds are frequently underestimated in the current literature.

The key antioxidant players, which catalyze the dismutation of O_2_^·-^ into H_2_O_2_, are superoxide dismutases (SODs), metalloenzymes utilizing metal cofactors such as nickel (NiSOD; not present in higher plants), manganese (MnSOD), iron (FeSOD) and zinc-copper (Cu/ZnSOD)^8^. The *Arabidopsis* genome encodes three Cu/ZnSODs (CSD1, CSD2, CSD3), one MnSOD (MSD1) and three FeSODs (FSD1, FSD2, FSD3) isoforms^9,10^.

The subcellular localization of individual SODs is linked to the detoxification requirements. MSD is responsible for scavenging of the superoxide generated in mitochondria^9^. FSD2 and CSD2 are reported to be attached to the thylakoid membrane of chloroplasts^11,12^, while FSD3 is colocalized with the chloroplast nucleoids and protects them against superoxide radicals through the formation of a heterodimeric protein complex with FSD2^12^. In turn, cytosolic localization is reported for two isoforms: CSD1 and FSD1^9,12^. Moreover, GFP-fusions suggest that FSD1 can localize to chloroplasts as well and deletion of the 11 amino-terminal nucleotides of FSD1 cDNA sequence restricted this protein to the cytosol^13^. However, the above mentioned studies relied on expression in either heterologous systems or protoplast cultures and there are currently no data on FSD1 *in vivo* localization *in planta*.

The absence or downregulation of some SODs cause phenotypic changes, suggesting their important roles in plant development. Knock-out *fsd2* and *fsd3* mutants display chlorotic phenotypes, abnormal chloroplast morphology and growth inhibition^12^. On the other hand, *fsd1* mutant does not show obvious phenotypes in green tissues or altered ROS levels in leaves when transferred into the dark for two days^12^. Nevertheless, overproduction of *Arabidopsis* FSD1 in *Zea mays* and *Nicotiana tabacum* caused increased tolerance against oxidative stress^14,15^. So far, root phenotypes of *fsd1* mutants have not been comprehensively studied. FSD1 protein shows high level of similarity with FSDs of agriculturally important crops such as *Brassica napus* (93% identity in amino acid sequence) or *Solanum lycopersicum* (75%) or *S. tuberosum* (74%), which is higher compared to *Arabidopsis* FSD2 (61%) and FSD3 (56%).

The major factor affecting FSD1 expression is the availability of copper in the culture medium, while Cu^2+^ homeostasis is mainly regulated by the transcription factor SQUAMOSA promoter binding protein-like 7 (SPL7)^16^. The expression of *SPL7* and *FSD1* genes during the day-light period is regulated by circadian and diurnal rhythms^17^. Furthermore, FSD1 activity is mediated by direct interaction with chloroplast chaperonin 20 (CNP20)^13^ and also by mitogen-activated protein kinases^18^.

In the present study, we aimed to gain new insights into the developmental expression and subcellular localization of FSD1 in *Arabidopsis* using advanced microscopy. We found that *FSD1* expression in living plants is tissue-specific and at the subcellular level, FSD1 localizes to the plastids, nuclei, and cytoplasm. Importantly, FSD1 was relocated to the plasma membrane after salt stress, which was correlated with periplasmic ROS production. Generally, our results provide new evidence for the specific localization and novel osmoprotective role for FSD1 in *Arabidopsis*.

## Materials and Methods

### Plant material and phenotyping

*Arabidopsis* seeds were surface sterilized by ethanol and placed on a 1/2 Murashige and Skoog (MS) medium solidified with 0.5% (w/v) gellan gum and stratified at 4°C for 1-2 days, to synchronize germination. For the preparation of 1/2 MS medium with different copper content, final CuSO_4_ · 5H_2_O concentrations were modified to 0 µM and 0.5 µM. Seedlings were grown vertically at 21°C, 16/8 h (light/dark) photoperiod with an illumination intensity of 150 μmol m^−2^ s^−1^ in a phytochamber (Weiss Technik, USA) for 1-15 days prior to imaging. For the preparation of etiolated plants, Petri plates were covered with aluminium foil.

*Arabidopsis* T-DNA knockout lines were obtained from the European Arabidopsis Stock Centre (http://arabidopsis.info/BasicForm; primers are listed in Supplementary Table 1). Two independent mutant lines *fsd1-1* (SALK_029455) and *fsd1-2* (GABI_740E11) were used, while the T-DNA insertion was confirmed by specific primers designed in the SIGnAL iSect tool (http://signal.salk.edu/tdnaprimers.2.html). Genomic DNA was isolated according to the manufacturer’s instructions of the Phire Plant Direct PCR Kit (Thermos Fisher Scientific, F130WH) and homozygous lines of mutants were confirmed by PCR.

For the detailed root phenotyping, seedlings were recorded daily and documented using a scanner (ImageScanner TM III, Little Chalfont, UK) and ZOOM stereo microscope (Axio Zoom.V16; Carl Zeiss, Germany) for two weeks. The primary root lengths of 7- and 10-day-old seedlings were measured from the individual scans in ImageJ (http://rsbweb.nih.gov/ij/). Lateral root number was counted on the 7^th^ and 10^th^ day after germination (DAG) and was standardized to the primary root length. The fresh weight of 14-day-old seedlings was measured. Phenotypic measurements were performed in three biological replicates (n=30) and the statistical significance was evaluated by one-way ANOVA test.

### Preparation of constructs and transgenic lines

Both C- and N-terminal fusion constructs of *eGFP* with genomic DNA of *FSD1* (*pFSD1-FSD1::GFP:3′UTR-FSD1* (GFP-FSD1) and *pFSD1::GFP:FSD1-3′UTR-FSD1* (FSD1-GFP)) were cloned under its native promoter from *Arabidopsis* wild type (Col-0). The sequence of the native promoter was taken 1270 bp upstream of the start codon and for 3′UTR 1070 bp downstream of the stop codon. MultiSite Gateway^®^ Three-Fragment Vector Construction (Thermo Fisher Scientific, 12537-023) was used as the cloning method for the preparation of these constructs. Amplified sequences of the promoter, genomic DNA and 3′UTR (primers are listed in Supplementary Table 1) were recombined into *pDONR™P4-P1R* and *pDONR™P2R-P3* donor vectors, where plasmids *pEN-L1-F-L2* with and without stop-codon were used as B fragment for the subsequent three-fragment vectors LR recombination into the destination vector *pB7m34GW*. Sequencing-validated cloning products were transformed into *Agrobacterium tumefaciens* GW3101, and used further for floral dip stable transformation of *fsd1-1* and *fsd1-2* mutants. Several transgenic lines possessing intense fluorescent signals have been selected from the T1 generation. Selected lines with one insertion were propagated into T3 homozygous generation and used in further experiments.

For immunoblotting analyses, a stably transformed A. thaliana G5 line expressing *35S:eGFP*^19^ was used as a positive control for GFP detection.

### Immunoblotting and SOD activity assay

Seedlings of each line were homogenized into fine powder in a mortar with liquid nitrogen. Proteins were extracted in E-buffer (50 mM HEPES pH 7.5, 75 mM NaCl, 1 mM EGTA, 1 mM MgCl_2_, 1 mM NaF, 10% (v/v) glycerol, PhosSTOP™ phosphatase inhibitor and Complete™ EDTA-free protease inhibitor coctail (both from Roche, Basel, Switzerland)) and the extract was centrifuged (13 000 g) at 4°C for 15 min. Protein concentrations of supernatants were measured using the Bradford assay. Equal amounts of proteins were mixed with 4-fold concentrated Laemmli Sample Buffer (Bio-Rad, Hercules, CA, USA) and boiled at 95°C for 5 min. Denatured protein extracts were separated by SDS-PAGE on 10% TGX Stain-Free™ Fast-Cast™ gels (Bio-Rad). Separated proteins were transferred to a polyvinylidene difluoride (PVDF) membrane (GE Healthcare, Little Chalfont, United Kingdom) using a wet tank unit (Bio-Rad) with Tris/glycine/methanol transfer buffer at 24 V and 4°C overnight. Nonspecific epitopes were blocked by overnight incubation of the membrane either in 5% (w/v) low-fat dry milk (for the detection of FSD1) or in 4% (w/v) low-fat dry milk and 4% (w/v) bovine serum albumin (for detection of GFP), both in Tris-buffered-saline with Tween 20 (TBS-T, 100 mM Tris-HCl; 150 mM NaCl; 0.1% Tween 20; pH 7.4). Subsequently, the membranes were incubated with anti-FSD primary antibody (Agrisera, dilution 1:3000 in TBS-T with 3% (w/v) low-fat dry milk) or anti-GFP (Sigma-Aldrich, dilution 1:1000 in TBS-T with 3% BSA) primary antibody at 4°C overnight. Following repeated washing in TBS-T, membranes were incubated with a secondary antibody diluted in TBS-T containing 1% (w/v) BSA for 1.5 h. Horseradish peroxidase-conjugated goat anti-rabbit and anti-mouse IgG secondary antibodies For the analysis of SOD isoenzymatic (both diluted 1:5000; Thermo Scientific) were used for the detection of FSD1 and GFP respectively. The signal was developed after five washing steps in TBS-T using the Clarity Western ECL substrate (Bio-Rad) and documented using the Chemidoc MP system (Bio-Rad).

For the analysis of SOD isoenzymatic activities, seedlings were homogenized in liquid nitrogen and subjected to protein extraction using 50 mM sodium phosphate buffer (pH 7.8), 1 mM ascorbate, 1 mM EDTA and 10% (v/v) glycerol. The extract was cleaned by centrifugation (13 000 g) at 4°C for 15 min, followed by measurement of the protein concentration. Samples of equal protein content were loaded on a 10% native PAGE gel and separated at constant 20 mA/gel for 2 h. Gels were preincubated in 50 mM sodium phosphate buffer, pH 7.8 for 10 min after separation. SOD isoform activities and their specific inhibition were visualized as described by Takáč et al. (2014)^18^.

The band intensities in immunoblots and native gels were quantified using Image Lab software (Bio-Rad). Both analyses were performed in three biological replicates and the statistical significance was evaluated using one-way ANOVA test.

### Quantitative analysis of transcript levels by quantitative real-time PCR

Isolation of total RNA from 14-day-old *Arabidopsis* seedlings (Col-0, *fsd1-1, fsd1-2* and GFP-FSD1 transgenic line) and subsequent quantitative real-time PCR (qRT-PCR) were performed according to Smékalová et al., 2014^20^. Experiments were run in three biological and three technical replicates. The expression data were normalized to the expression of elongation factor 1-alpha (EF1α) used as a reference gene (primers are listed in Supplementary Table 1). Statistical significance was tested by one-way ANOVA test.

### Whole mount immunofluorescence labelling

*Arabidopsis* Col-0 and *fsd1* mutants grown on 1/2 MS medium were used at 3^rd^ DAG for immunofluorescence labeling of the root tips according to the protocol established by Šamajová et al. (2014)^21^ with minor modifications. Samples were incubated with rat anti-FSD1 (Agrisera) primary antibody diluted at 1:250, in phosphate-buffered saline (PBS) containing 3% (w/v) BSA at 4°C overnight. In the next step, samples were incubated with Alexa-Fluor 488 conjugated goat anti-rat secondary antibody diluted at 1:500 in PBS with 3% (w/v) BSA at room temperature for 3 h. DNA was counterstained with 250 μg/ml 4,6-diamidino-2-phenylindole (DAPI, Sigma-Aldrich) in PBS for 10 min. After a final wash in PBS, the specimens were mounted in an antifade solution (0.5% (w/v) p-phenylenediamine in 70% (v/v) glycerol in PBS or 1 M Tris-HCl, pH 8.0) or in the commercial antifade Vectashield™ (Vector Laboratories).

### Salt sensitivity assay and plasmolysis

Germination analysis of Col-0, both *fsd1* mutants and *fsd1-1* complemented lines (GFP-FSD1 and FSD1-GFP) was performed on ½ MS medium with and without 150 mM NaCl. Plates with seeds were kept at 4°C for 2 days and incubated as mentioned above. Percentage of germinated seeds (with visible radicle) was counted under stereomicroscope after 24, 48, and 78 hours. Measurements were performed in four repetitions (n=30) and statistical significance was tested by one-way ANOVA test.

For salt stress sensitivity determination, 4-day-old seedlings of Col-0, *fsd1* mutants and *fsd1-1* complemented lines (GFP-FSD1 and FSD1-GFP) growing on ½ MS medium were transplanted to ½ MS medium containing 150 mM NaCl. The ratio of bleached seedlings was counted at the 5^th^ day after transfer. Measurements were performed in four repetitions (n=30) and the statistical significance was evaluated by one-way ANOVA test.

For plasmolysis induction, 4-day-old seedlings of *fsd1-1* complemented lines (FSD1-GFP and GFP-FSD1) were mounted between glass slide and coverslip in liquid 1/2 MS media. Plasmolysis was induced with 500 mM NaCl (hypocotyls) or 250 mM NaCl (roots) in liquid 1/2 MS media applied by perfusion. Plasmolyzed cells were observed 5-30 min after the perfusion by CLSM 880 equipped with an Airyscan detector (ACLSM, Carl Zeiss, Germany) and a spinning disk microscope (Cell Observer, SD, Carl Zeiss, Germany).

### Histochemical and fluorescent detection of ROS

To visualize superoxide production in roots, 7-day-old seedlings of Col-0, *fsd1* mutants and *fsd1-1* complemented lines were incubated in 10 mM potassium phosphate buffer (pH 7.8) containing 0.02% (w/v) 4-nitroblue tetrazolium chloride (NBT) for 5 min in dark. Stained seedlings were boiled in clearing solution containing 20% (v/v) acetic acid, 20% (v/v) glycerol and 60% (v/v) ethanol for 5 min and stored in mixture of 20% glycerol (v/v) and 80% (v/v) ethanol. Reduced NBT was visualized as a dark blue-colored formazan deposit.

ROS in plasmolyzed roots were visualized by incubation in 30 µM 2’,7’-dichlorodihydrofluorescein diacetate (CM-H_2_DCFDA), diluted in ½ MS with or without 250 mM NaCl for 15 min in darkness. The emitted signal (excited at 492-495 nm) was recorded at 517-527 nm using CLSM 720 (Carl Zeiss, Germany).

### Confocal laser scanning microscopy

Seedlings of *fsd1-1* mutants carrying recombinant GFP-fused FSD1 were used for microscopy at 3^th^−8^th^ DAG. Imaging of living or fixed samples was performed using a confocal laser scanning microscope LSM710 (Carl Zeiss, Germany), LSM880 equipped with an Airyscan (ACLSM, Carl Zeiss, Germany) and a spinning disk microscope (Cell Observer, SD, Carl Zeiss, Germany). Image acquisition was done with 20× (0.8 NA) dry Plan-Apochromat, 40× (1.4 NA) and 63× (1.4 NA) Plan-Apochromat oil-immersion objectives. Samples were imaged with a 488 nm excitation laser using emission filters BP420-480+BP495-550 for eGFP detection and BP 420-480 + LP 605 for chlorophyll *a* detection. Laser excitation intensity did not exceed 2% of the available laser intensity range. Immunolabelled samples were imaged using the excitation laser line 488 nm and emission spectrum 493-630 nm for Alexa-Fluor 488 fluorescence detection, and excitation laser line 405 nm and emission spectrum 410-495 nm for DAPI. Living plants of 3^th^-8^th^ DAG were stained with 4 μM FM4-64 (Invitrogen, USA) diluted in 1/2 liquid MS medium for 10 min before imaging. Samples were observed with excitation laser line 488 nm for eGFP detection and 561 nm for FM4-64 detection. Images were processed as maximum intensity projections of Z-stacks in Zen Blue 2012 software (Carl Zeiss, Jena, Germany), assembled and finalized in Microsoft PowerPoint to final figures.

### Light-sheet fluorescence microscopy

Seeds of *fsd1-1* mutant expressing *proFSD1::FSD1:GFP* constructs were surface-sterilized and placed on 1/2 MS medium solidified with 0.5% (w/v) gellan gum and stratified at 4°C for 1-2 days. Subsequently, seeds were transferred to horizontally-oriented plates with the same culture medium and a height of at least 15 mm. Horizontal cultivation allowed seeds to germinate and roots to grow inside of a solidified medium. Seedlings were inserted into fluorinated ethylene propylene (FEP) tubes with an inner diameter of 2.8 mm and wall thickness of 0.2 mm (Wolf-Technik, Germany), in which roots grew in the block of the culture medium inside the FEP tube, while the upper green part of the seedling developed in an open space of the FEP tube with access to the air^22^. The FEP tube with seedling was inserted into a sample holder and placed into the observation chamber of the light-sheet Z.1 fluorescence microscope (Carl Zeiss, Germany). Before insertion of the sample into the microscope, plants were ejected slightly out of the FEP tube allowing for imaging of the root in the block of the solidified culture medium, but without the FEP tube. The sample chamber of the microscope was filled with sterile 1/2 MS medium and tempered to 22°C using the peltier heating/cooling system.

Developmental live cell imaging was done with dual-side light-sheet illumination using two LSFM 10x/0.2 NA illumination objectives (Carl Zeiss, Germany) with excitation laser line 488 nm, beam splitter LP 560 and with emission filter BP505-545. Image acquisition was done with a W Plan-Apochromat 20x/1.0 NA objective (Carl Zeiss, Germany) and images were recorded with the PCO. Edge sCMOS camera (PCO AG, Germany) with an exposure time of 100 ms and imaging frequency of every 2 min in the Z-stack mode for 2-20 hours.

## Results

### FSD1 is developmentally regulated in the early post-germination phase of plant growth

According to the public expression data deposited in the Genevestigator database^10^, *FSD1* is developmentally regulated and is abundantly expressed at early developmental stages. Generally, *FSD1* expression prevails at the vegetative growth phase, while *CSD1, CSD2* and *MSD1* isoforms are typically expressed during the reproductive phase^10^. Analysis of FSD1 abundance and activity during *Arabidopsis* early seedling growth revealed that both parameters gradually increased from the 3^rd^ to 13^th^ DAG, but significantly decreased in following days (Fig. 1a-d). In order to address the possible phenotypic consequences of FSD1 deficiency at early developmental stages, two independent homozygous T-DNA insertion *fsd1* mutants were analyzed. It was found that both mutants exhibited reduced lateral root density, while no significant difference was found in the primary root length and seedling fresh weight compared to the wild type (Fig. 1e-h). In summary, our data suggest that, FSD1 activity and abundance in *Arabidopsis* depends on the growth phase and its deficiency leads to reduced lateral root numbers.

**Fig. 1.**
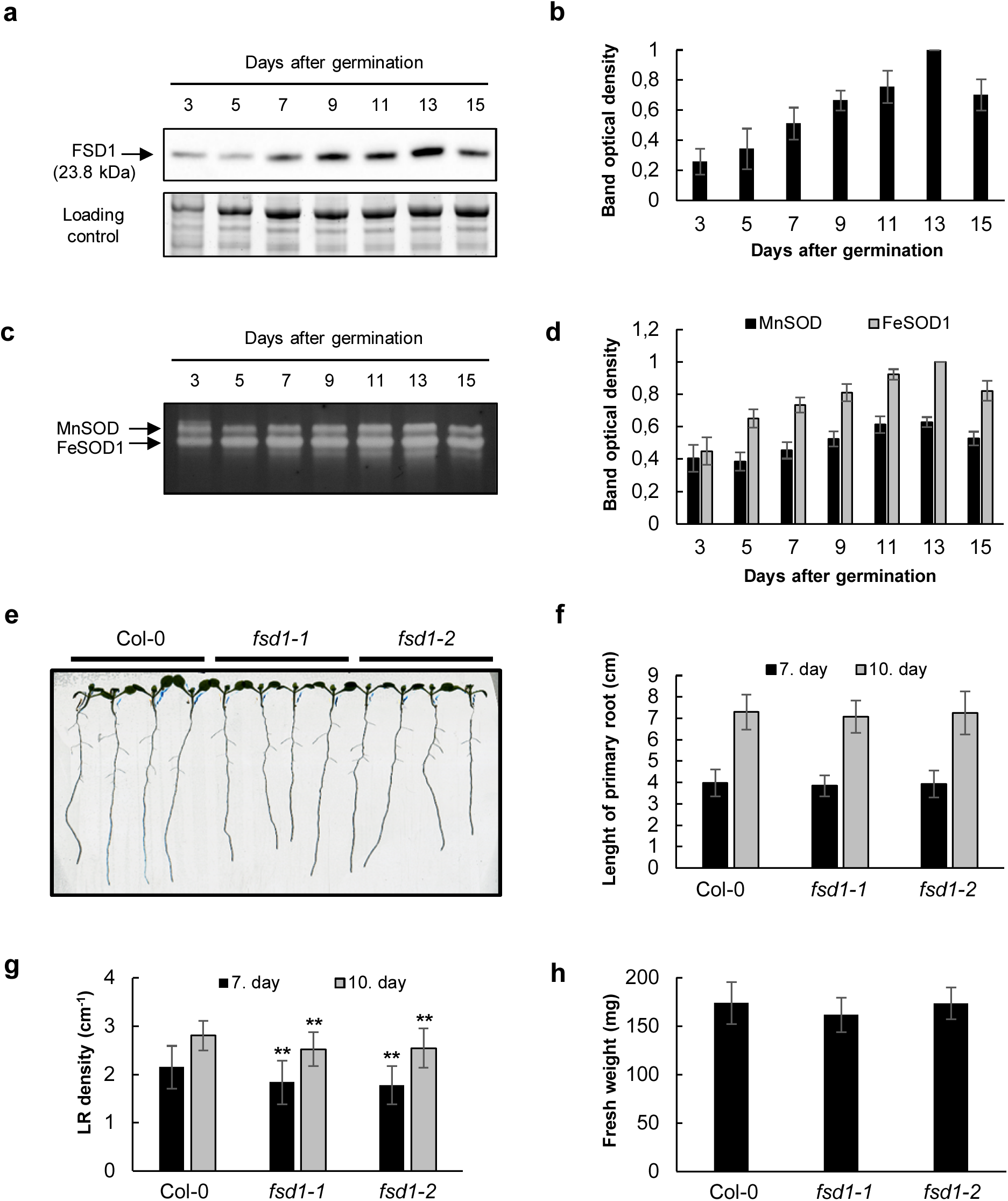
Early developmental and phenotypical analysis of iron superoxide dismutase 1 (FSD1). **a**, Immunoblotting analysis of FSD1 abundance using anti-FSD1 antibody during early development of *Arabidopsis* (Col-0) seedlings. **b**, Quantification of optical densities of bands in **(a)**. The densities are expressed as relative to the highest value. **c**, Visualization of SOD isoform activities on native polyacrylamide gels during early development of *Arabidopsis* wild type (Col-0) seedlings. **d**, Quantification of optical densities of bands in **(c)**. The densities are expressed as relative to the highest value. **e**, Representative image of *fsd1-1* and *fsd1-2* mutant and Col-0 seedlings on 7^th^ day after germination (DAG). **f-h**, Quantification of primary root length **(f)**, lateral root density **(g)** of indicated seedlings on 7^th^ and 10^th^ DAG and fresh weight of seedlings on 14^th^ DAG **(h)**. Error bars represent standard deviation. Stars indicate statistically significant difference as compared to Col-0 (one-way ANOVA, **p < 0.01).

### Functional complementation of *fsd1* mutants

For the elucidation of FSD1 expression and localization *in vivo*, we generated stably transformed *fsd1* mutants carrying FSD1 under its own native promoter and fused with GFP. Both N- and C-terminal GFP fusions were cloned and individually introduced into *fsd1* mutants. FSD1 complementation reverted the lateral root phenotypes of *fsd1* mutants (Fig. 2a). In addition, primary root length (Fig. 2b), lateral root density (Fig. 2c), and seedling fresh weight (Fig. 2d) in complemented lines slightly exceeded the respective values in wild-type plants. Neither FSD1 protein presence, nor enzymatic activity were observed in *fsd1* mutants by biochemical analyses (Fig. 2e-h), while GFP-tagged FSD1 proteins (FSD1-GFP or GFP-FSD1) were detected in both complemented lines (Fig. 2e-h, Supplementary Fig. 1). Quantitatively, wild type-like level of FSD1 activity and abundance was found in FSD1-GFP complemented plants, as examined by both anti-FSD (Fig. 2e, f) and anti-GFP antibodies (Supplementary Fig. 1). On the other hand, strongly reduced (representing 70% and 56% of wild type as examined by anti-FSD and anti-GFP antibodies, respectively) protein levels were found in the GFP-FSD1 complemented line (Fig. 2e, f, and Supplementary Fig. 1). Quantitative PCR analysis showed that *FSD1* transcript levels were similar to wild type (Supplementary Fig. 2). Functionality of the FSD1 proteins fused with GFP in both complemented lines was shown by the detection of their activities (Fig. 2g, h). Moreover, FSD1 activities and abundances of both GFP-FSD1 and FSD1-GFP were sensitive to copper content in cultivation media, further confirming their functionality (Supplementary Fig. 3).

**Fig. 2.**
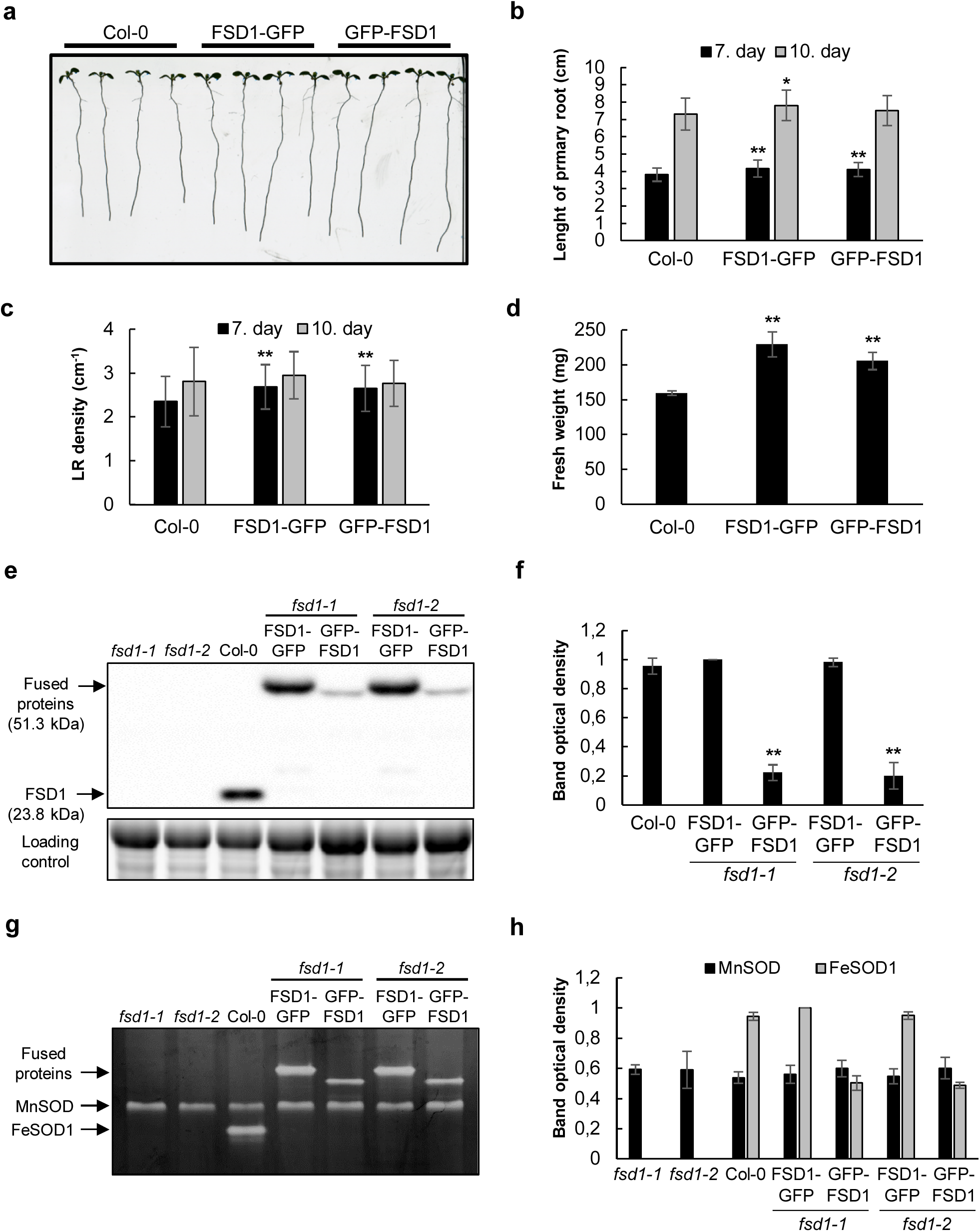
Phenotypic and functional analysis of *fsd1* complemented mutants. **a**, Representative image of 7-day-old Arabidopsis wild type (Col-0) and *fsd1-1* mutant seedlings expressing *proFSD1::FSD1:GFP* or *proFSD1::GFP:FSD1*. **b,c**, Quantification of primary root length **(b)** and lateral root density **(c)** of indicated 7- and 10-day-old seedlings. **d**, Fresh weight of indicated 14-day-old seedlings. **e**, Immunoblotting analysis of FSD1, FSD1-GFP and GFP-FSD1 abundance in 14-day-old *fsd1* mutants, Col-0 and complemented *fsd1* mutants using anti-FSD antibody. **f**, Quantification of band optical densities in **(e)**. The densities are expressed as relative to the highest value. **g**, Visualization of activities of SOD isoforms on native polyacrylamide gels in indicated plant lines. **h**, Quantification of optical densities of bands in **(g)**. The densities are expressed as relative to the highest value. Error bars represent standard deviation. Stars indicate statistically significant difference as compared to Col-0 (one-way ANOVA, *p < 0.05, **p < 0.01).

Inhibition of FSD1 activity by H_2_O_2_, but not by KCN suggests that the bands on the native PAGE gels correspond to FSD1 proteins fused with GFP (Supplementary Fig. 4). FSD1 activities in the transgenic lines quantitatively correlate with the abundances of the respective fused and native proteins (Fig. 2g, h and Supplementary Fig. 4a). Interestingly, the band corresponding to GFP-FSD1, migrated in a distinct manner as compared to FSD1-GFP on the native PAGE gel (Fig. 2g). We also tested whether these manipulations with FSD1 expression resulted in different endogenous levels of superoxide. The histochemical examination of superoxide in mutant and transgenic lines did not show any differences in comparison to the wild type or among the lines (Supplementary Fig. 5).

Together, these results suggest that FSD1 is important for the fine-tuning of the lateral root development.

### Expression pattern of FSD1-GFP during germination and early seedling development

Spatial and temporal patterns of FSD1-GFP expression in the early stages of development were monitored *in vivo* using light-sheet fluorescence microscopy. This allowed the time-lapse monitoring of FSD1-GFP distribution during the whole process of seed germination at nearly environmental conditions (Fig. 3, Supplementary Video 1). Within the first 6 h of seed germination, still before radicle emergence, we observed an increase of FSD1-GFP signal in the micropylar endosperm with a maximum at the future site of radicle protrusion (Fig. 3a-g, Supplementary Video 1). With the endosperm rupture and emergence of the primary root, FSD1-GFP signal gradually decreased in the micropylar endosperm (Fig. 3h-j), while a strong FSD1-GFP signal appeared in the fast-growing primary root (Fig. 3k, l, Supplementary Video 1). Strong expression of FSD1-GFP was visualized in the transition and elongation zones of the primary root (Fig. 3l, m, Supplementary Video 1), which was, however, gradually decreasing in the differentiation zone, particularly after the emergence of the root hairs in the collar region (Fig. 3m-o). During seed germination, FSD1-GFP-labelled plastids in endosperm cells showed a high degree of motility (Supplementary Video 1). Thus, FSD1 may be involved in the process of endosperm rupture during seed germination. Moreover, FSD1 tissue-specific expression might play a protective role during early root emergence from seeds.

**Fig. 3.**
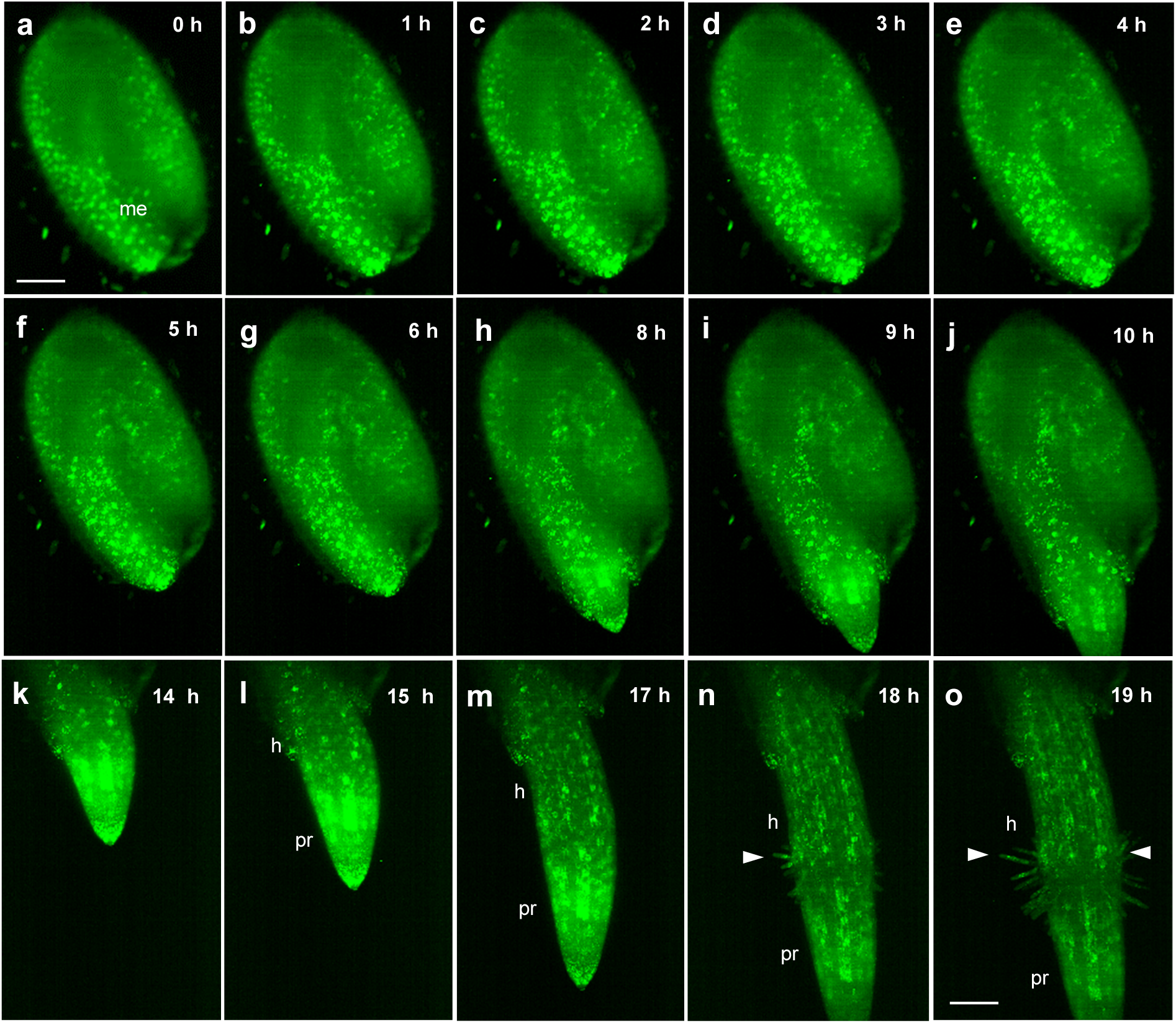
Time-lapse monitoring of FSD1-GFP distribution during seed germination obtained using light sheet fluorescence microscopy. **a-e**, sequential accumulation and relocation of the signal in micropylar endosperm (me) to the site of radicula protrusion. **e**, testa rupture. **f-h**, radicula protrusion. **h**, endosperm rupture. **k-o**, primary root elongation. **n,o**, primary root differentiation. Arrowheads point to the site of root hairs in the collar region on the border between the elongating primary root (pr) and hypocotyl (h). Scale bar: 100 µm.

After germination, which occurred during the 1^st^ DAG, growth of the primary root continued and cotyledons were released from the seed coat during the 2^nd^ DAG (Supplementary Fig. 6a, b). Expression levels of FSD1-GFP in emerging cotyledons were high (Supplementary Fig. 6b). Hypocotyl and fully opened cotyledons in developing seedlings at 5^th^ DAG contained moderate amount of FSD1-GFP, while the strongest signal was detectable in the shoot apex and emerging first true leaves (Supplementary Fig. 6c). FSD1-GFP signal considerably increased in the lateral root primordia (Supplementary Fig. 6d-f). Accumulation of FSD1-GFP was still visible in the apices of the lateral roots as well as in the basal parts, at the connection of the lateral roots to the primary root (Supplementary Fig. 6g). In growing apex of the primary root, the strongest FSD1-GFP signal was located in the transition zone (Supplementary Fig. 6h). The FSD1-GFP signal gradually decreased with acceleration of the cell elongation, differentiation, and root hair formation (Supplementary Fig. 6h, Supplementary Video 2).

### Tissue-specific subcellular localization of GFP-FSD1 and FSD1-GFP in *Arabidopsis*

In the cells of both above- and underground organs of light-exposed seedlings of *fsd1-1* mutants harboring *proFSD1::FSD1:GFP* construct, C-terminal FSD1-GFP fusion protein was localized in plastids, nuclei, and cytoplasm, especially in the cortical cytoplasmic layer in close proximity to the plasma membrane (Supplementary Videos 3 and 4). Such localization patterns of FSD1-GFP was consistent in cells of all aboveground organs in light-exposed seedlings, such as cotyledon epidermis (mature pavement cells, stomata and their precursors, Fig. 4a-c; Supplementary Videos 3 and 4), leaf mesophyll cells (Fig. 4d-f, Supplementary Video 5), hypocotyl epidermis (Fig. 4g), and first true leaf epidermis with branched trichomes (Fig. 4h). In leaf pavement cells, FSD1-GFP-labelled plastids were located around the nucleus and in the cytoplasmic strains traversing the vacuole (Fig. 4a). Plastids located in cytoplasmic strands actively followed other organelles during cyclosis (Supplementary Video 3 and 4). Some other FSD1-GFP-labelled plastids located in a close proximity to nuclei in stomata guard cells and adjacent pavement cells, were less dynamic (Supplementary Videos 3 and 4). In mesophyll cells, FSD1-GFP-labelled plastids were temporarily contacted and eventually interconnected by the highly dynamic network of tubules and cisternae of the endoplasmic reticulum (Supplementary Video 5).

**Fig. 4.**
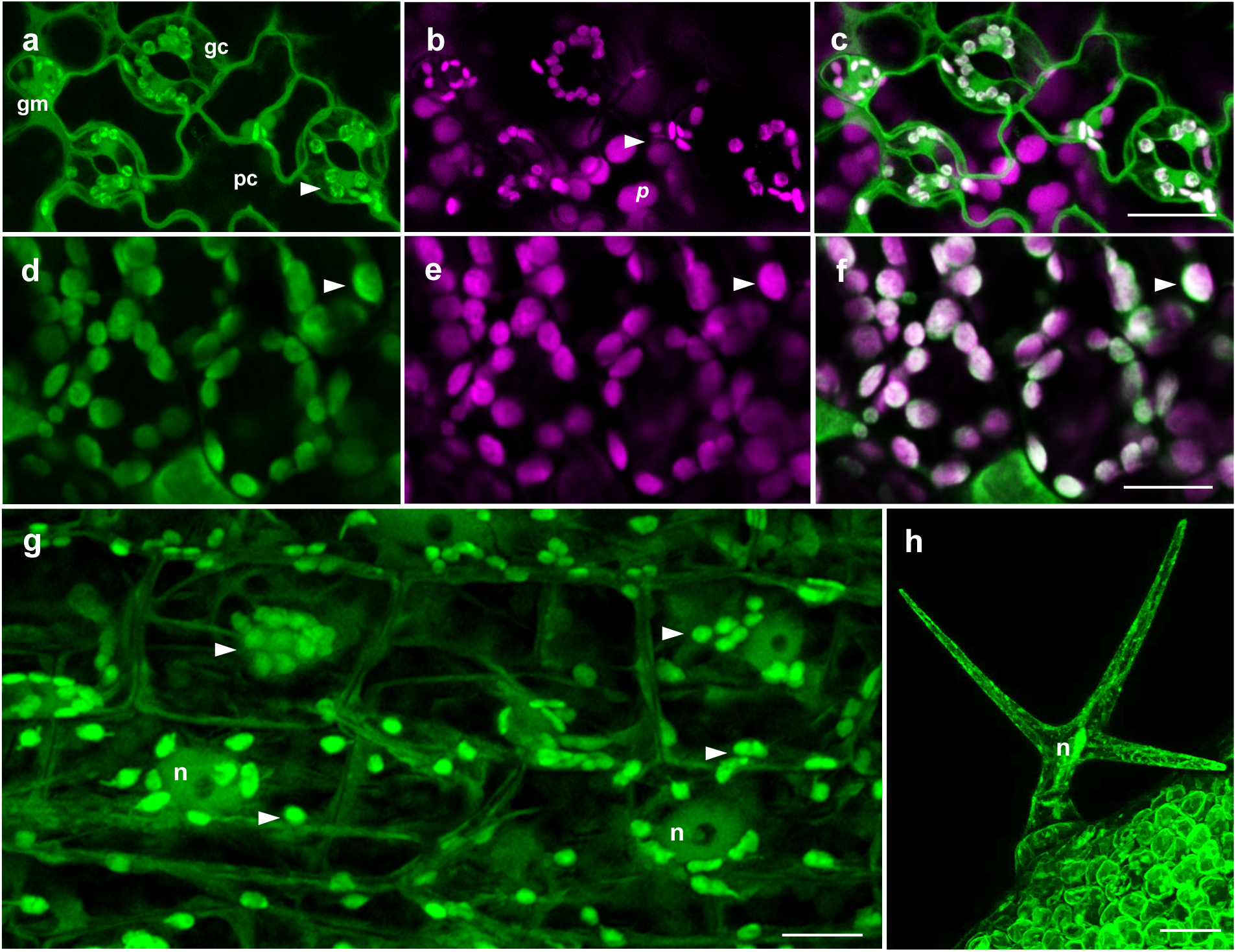
FSD1-GFP localization in cells of *Arabidopsis* aboveground organs revealed by Airyscan confocal laser scanning microscopy. **a-c**, adaxial surface of cotyledons with pavement (*pc*), guard (*gc*) and guard mother (*gm*) cells. **d-f**, mesophyll cells. **g**, epidermal cells of hypocotyls. **h**, triple-branched leaf trichome. Indications: (*n*) nucleus. Arrowheads point on accumulation of FSD1-GFP in plastids. Channels: green - FSD1-GFP; magenta - chlorophyll *a* autofluorescence. Scale bars: **a-g**, 10 μm; **h**, 20 μm.

Moreover, FSD1-GFP maintained the same pattern of its localization in cotyledon epidermal cells of etiolated *Arabidopsis* seedlings, although it was more intensively accumulated in the cortical cytoplasm just beneath the plasma membrane as compared to the light-exposed plants (Supplementary Fig. 7). In turn, FSD1-GFP was abundant in etioplasts, showing only basal remaining level of chlorophyll *a* autofluorescence (Supplementary Fig. 7b, c).

Plastidial, nuclear and cytoplasmic localization of FSD1-GFP was detected also in cells of the root apex (Fig. 5a, Supplementary Video 6). This localization pattern was visible in cells of the lateral root cap (Fig. 5a, b,), in meristematic cells (Fig. 5a, c), epidermal cells of elongation zone (Fig. 5d, e) as well as in trichoblasts within the differentiation zone (Fig. 5f) of primary root. The selective styryl dye FM4-64 counterstaining of the plasma membrane in root cells helped to reveal tissue-specific FSD1-GFP localization in the root tip (Supplementary Fig. 8). It showed lower GFP-FSD1 signal intensity in central columella cells (Fig. 5a, Supplementary Fig. 8).

**Fig. 5.**
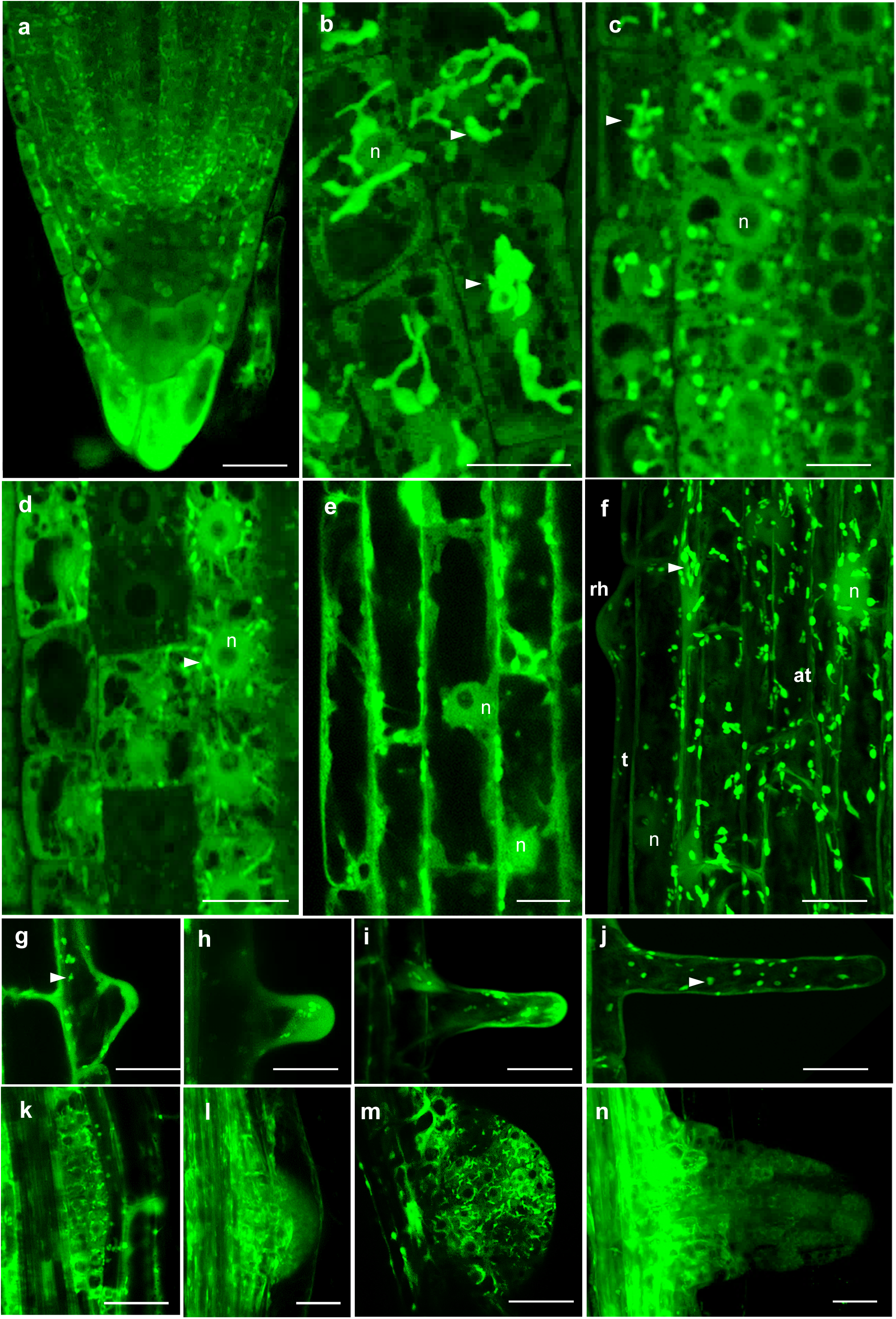
Tissue- and organ-specific subcellular FSD1-GFP localization in *Arabidopsis* roots revealed by Airyscan confocal laser scanning microscopy. **a**, primary root apex. **b**, root cap cells with GFP-signal in plastids (arrowheads) and nuclei (n). **c**, epidermal and cortical meristem cells. **d**, cortical cells of distal elongation zone. **e**, cortical cells of elongation zone. **f**, trichoblasts (t) with an emerging root hair (rh) and atrichoblasts (at) of differentiation zone. **g-j**, mid-plane sections of root hairs. **g**, primordia. **h, i**, elongating root hair. **j**, mature root hair. **k-m**, mid-plane sections of forming lateral root primordia at diverse developmental stages, 4^th^ day after germination (DAG). **o**, emerged lateral root, 8^th^ DAG. Scale bars: **a, e, f, k-n**, 20 μm; **b, c, d, g-j** 10 μm.

Furthermore, accumulation of FSD1-GFP was observed in the lateral root primordia emerging from the pericycle (Fig. 5k-n). FSD1-GFP signal increased first in cells of forming lateral root primordium still enclosed by tissues of the primary root (Fig. 5k, Supplementary Fig. 9a-c). Strong signal of FSD1-GFP was found in cells of the central region, where the apical meristem of the emerging lateral root was established (Fig. 5l, m). Considerably high levels of FSD1-GFP also persisted during the release of the lateral root from the primary root tissue (Fig. 5n, Supplementary Fig. 9d-f). Established apex of elongating lateral root showed differential pattern of FSD1-GFP expression, with high levels in the endodermis/cortex initials (Supplementary Fig. 9g-i, Supplementary Video), actively dividing cells of the epidermis, cortex and endodermis, and lateral root cap cells (Supplementary Fig. 9g-i). On the other hand, considerably lower levels of FSD1-GFP occurred in cells of the quiescence center and columella (Supplementary Fig. 9g-i).

The process of root hair formation from trichoblasts was connected with the accumulation of FSD1-GFP in the cortical cytoplasm of the emerging bulge (Fig. 5g). In tip-growing root hairs, FSD1-GFP accumulated in the apical and subapical zone (Fig. 5h, i). It is noteworthy that after the termination of root hair elongation, FSD1-GFP signal (Fig. 5j) dropped at the tip, while typical strong plastidial signal appeared in the cortical cytoplasm (Fig. 5j).

Subcellular localization pattern of FSD1 was confirmed by the whole mount immunofluorescence localization method in fixed samples using anti-FSD antibody. This technique showed prominent strong immunolocalization of FSD1 to plastids distributed around nuclei and in the cytoplasm, as well as nuclear and cytoplasmic localization in meristematic cells of the primary root (Fig. 6a-f).

**Fig. 6.**
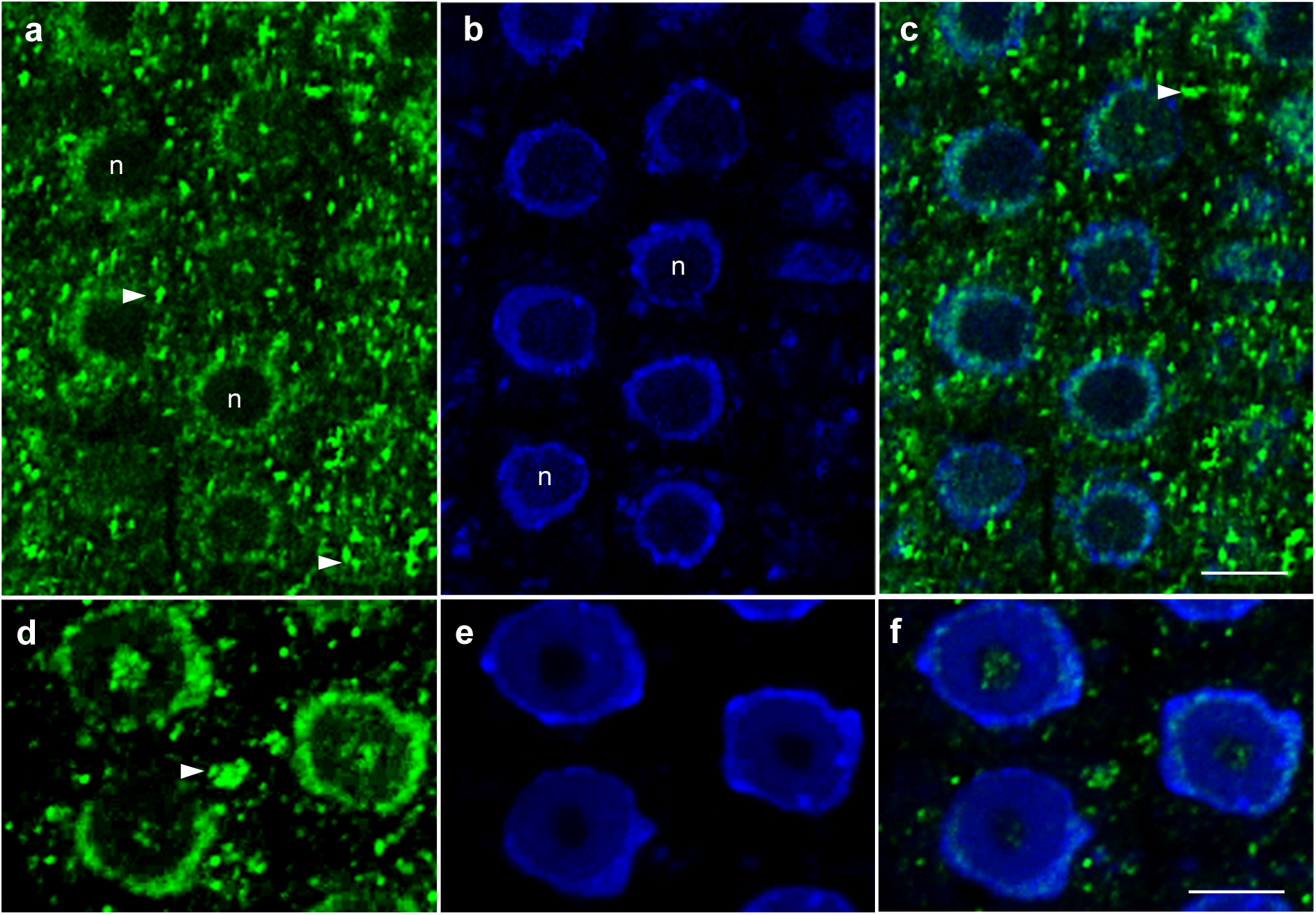
Overview of FSD1 immunolocalization in interphase meristematic cells of *Arabidopsis* (Col-0) primary roots. The images represent maximum intensity projections of 20 optical sections (with thickness of 0.18 µm each) at the mid-plane of root meristem cells with **a-c** or without **d-f** deconvolution in ZEN Blue 2012 software. Green immunolabelling with anti-FeSOD - Alexa Fluor 488); blue - DAPI staining. Arrowheads indicate plastids. (n) stands for nuclei. Scale bar: 5 µM.

Interestingly, the N-terminal GFP-FSD1 fusion protein was not targeted to plastids, but it was localized both in the nuclei and cytoplasm. This localization pattern was observed in leaf pavement (Supplementary Video 7) and stomata guard cells (Supplementary Fig. 10a-c), in cotyledon mesophyll cells (Supplementary Fig. 10d-f) as well as in hypocotyl epidermal cells (Supplementary Fig. 10g, Supplementary Video 8). The absence of plastidial localization did not affect the tissue-specific expression pattern of GFP-FSD1 in primary root apex. The strongest signal was located in the epidermis, cortex, endodermis, and root cap (Supplementary Fig. 10h). Considerably lower GFP-FSD1 signal was detected in the quiescent center, central columella cells and proliferating tissues of the central cylinder (Supplementary Fig. 10h). Strong accumulation of GFP-FSD1 was typically present in founding cells of the lateral root primordia and adjacent pericycle cells (Supplementary Fig. 10i). Taking into account the strong reduction in FSD1 abundance and activity in transgenic line expressing N-terminal GFP-FSD1 fusion as compared to FSD1-GFP (Fig. 2e-h, Supplementary Fig. 1), the plastidic FSD1 pool may represent around half of the total FSD1 pool in *Arabidopsis* cells. Notably, the level of *FSD1* transcripts in GFP-FSD1 line was comparable to the wild-type transcript level (Supplementary Fig. 2), indicating possible degradation of plastid-targeted FSD1 in the GFP-FSD1 line.

Plastids were the organelles most strongly accumulating FSD1-GFP and located either around the nuclei or distributed throughout the cytoplasm (Fig. 4, Fig. 5, Supplementary Fig. 7, Supplementary Video 3 and 4). Typically, plastids in cells of different tissues formed polymorphic stromules, which displayed different tissue-specific shape, length, branching (Fig. 4, Fig. 5) and dynamicity (Supplementary Videos 3,5). Thus, in lateral root cap cells highly dynamic FSD1-GFP-labelled plastids persistently formed long stromules, touching each other (Fig. 5b, Supplementary Video 9), while the plastids in isodiametric meristematic cells possessed less stromules (Fig. 5c, d). In hypocotyl epidermal cells with active cytoplasmic streaming, only some plastids were interconnected by stromules (Supplementary Video 10). Since stromules are tubular plastid extensions filled with stroma^23^, FSD1 might be considered a stromal protein. In contrast to FSD2 and FSD3^12^, FSD1 was not detected in the chloroplast nucleoids.

### FSD1 contributes to salt stress tolerance in *Arabidopsis* by superoxide conversion in the periplasm

Protective role of FSD1 during the early stages of post-embryonic plant development was tested in *fsd1* mutants and complemented lines on seed germination under salt stress conditions. Seed germination of *fsd1* mutants was strongly reduced by the presence of 150 mM NaCl in the 1/2 MS medium, while FSD1-GFP lines exhibited germination rates comparable to that of wild type. GFP-FSD1 line showed an insignificantly reduced germination rate on the 1^st^ day, but germination efficiency was synchronized with wild type and FSD1-GFP line from the 2^nd^ day onwards (Fig. 7a). The results indicated that FSD1 expressed under its own native promotor functionally complemented the salt stress-related deficiency of *fsd1* mutants.

**Fig. 7.**
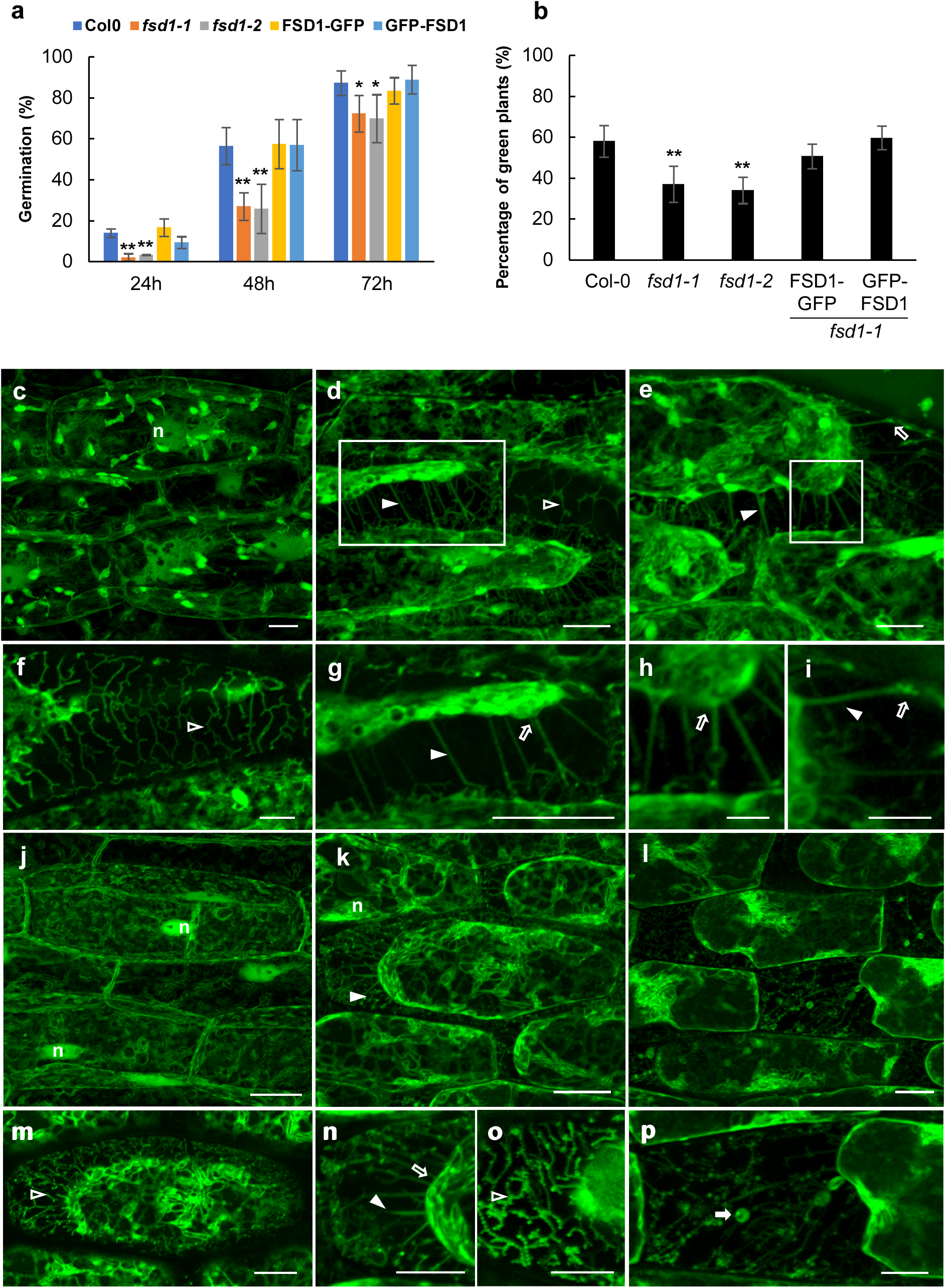
Response of *fsd1* mutants and complemented mutant lines to salt stress. **a**, Seed germination efficiency on 150 mM NaCl. **b**, Viability of seedlings on 4^th^ day after the transfer to 150mM NaCl-containing medium. Stars indicate statistically significant difference as compared to Col-0 (one-way ANOVA, *p < 0.05, **p < 0.01). **c-i**, FSD1-GFP signal in hypocotyl epidermal cells on ½ MS **(c)** and 500 mM NaCl (15 min) **(d-i)**. Images showing Hechtian reticulum **(f)** and strands **(g)** are close-ups from image **(d). h-i**, Hechtian strands and their connections to plasma membrane, close-ups from **(e). j-p**, GFP-FSD1 in hypocotyl epidermal cells exposed to ½ MS **(j)** and 500 mM NaCl **(k-p)** for 15 min. Hechtian reticulum **(m)** and strands **(o)** are close-ups from **(k)**. Disturbed Hechtian reticulum with aggregations **(p)** is close-up from **(l)**. Filled arrowheads indicate Hechtian strands; blank arrowheads - Hechtian reticulum; filled arrows - globular aggregations; blank arrows - connections of Hechtian strands to plasma membrane and cell wall. Scale bar: **a-g, j-p**, 10 µm; **h,i**, 5 µm.

To further test the new role of FSD1 in salt stress sensitivity, we characterized the response of developing seedlings to the high salt concentration in the culture medium. We found that both *fsd1* mutants showed hypersensitivity to NaCl and exhibited increased cotyledon bleaching. Both FSD1-GFP and GFP-FSD1 fusion proteins efficiently reverted the salt hypersensitivity of *fsd1* mutants (Fig. 7b, Supplementary Fig. 11). These results supported the new functional role of FSD1 in *Arabidopsis* salt stress tolerance.

To gain deeper insight into FSD1 function during plant response to the salt stress, we performed subcellular localization of FSD1-GFP in hypocotyl epidermal cells plasmolyzed by 500 mM NaCl (Fig. 7c-i, Supplementary Fig. 12). In addition to plastidial, nuclear, and cytoplasmic localization in untreated cells (Fig. 7c), FSD1-GFP was detected in Hechtian strands and Hechtian reticulum, interconnecting retracted protoplast with the cell wall of plasmolyzed cells (Fig. 7d-i). Hechtian reticulum located in close proximity to the cell wall (Fig. 7f), and thin attachments of Hechtian strands to peripheral Hechtian reticulum in the form of bright spots (Fig. 7e, g-i) were enriched with FSD1-GFP (Fig. 7h, i, Supplementary Fig. 12). Plasmolyzed cells showed strong GFP signal at plasma membrane and also contained vesicle-like structures decorated by FSD1-GFP, in their cytoplasm (Supplementary Fig. 12d) and also within the Hechtian strands (Fig. 7h). We observed a similar relocation pattern in the GFP-FSD1 line. GFP-FSD1 was located in the nuclei and cytoplasm of untreated cells (Fig. 7j), while prominent GFP-FSD1 accumulation was observed at the plasma membrane of retracted protoplasts, in Hechtian strands and reticulum after plasmolysis (Fig. 7k-p). Peripheral Hechtian reticulum and strands were decorated by spot- and vesicle-like structures labelled with GFP-FSD1 (Fig. 7l, p).

Subcellular localization during the plasmolysis induced by salt stress implies that FSD1 could be involved in the ROS production within the periplasmic space. Therefore, we used a fluorescent ROS indicator CM-H_2_DCFDA, which is preferentially specific to H_2_O_2_^24^, for intracellular ROS localization in plasmolyzed cells. We have found that the CM-H_2_DCFDA fluorescence signal highly correlated with the subcellular distribution of GFP-tagged FSD1 after plasmolysis (Fig. 8a-f; Supplementary Fig. 13a, b). Intense ROS production was detected in the cytoplasm of retracted protoplasts (Fig. 8a), as well as in Hechtian strands and reticulum connecting them to the cell walls (Fig. 8a). Interestingly, branched Hechtian reticulum (Fig. 8a, d), vesicular-like structures within Hechtian strands (Fig. 8a-c) and connecting points of Hechtian strands to the cell wall (Fig. 8a, f) were the places of intense ROS production. Collectively, these data indicate that at least some part of salt stress-induced ROS production and accumulation at the plasma membrane and Hechtian strands and reticulum depends on relocated FSD1.

**Fig. 8.**
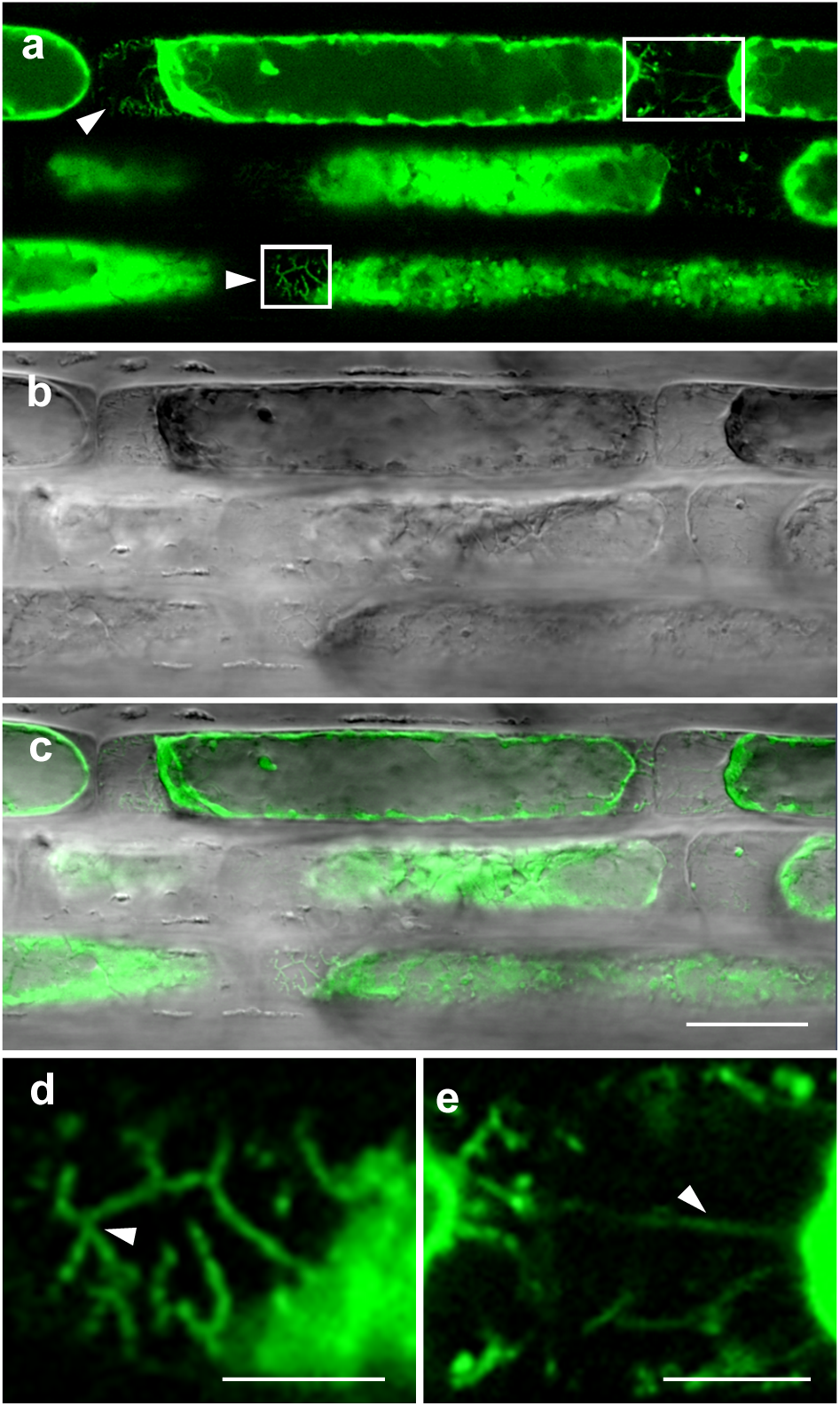
Accumulation of reactive oxygen species (ROS) in *Arabidopsis* primary root in response to salt stress. Plasmolysis was induced by the treatment of 4-day-old seedlings with liquid ½ MS medium containing 250 mM NaCl for 15 min. **a-e**, ROS distribution during the plasmo**e**lysis visualized by fluorescent tracker CM-H_2_DCFDA. **b**, transmitted light. **c**, overlay. **d, e**, details of ROS accumulation on Hechtian strains and reticulum (arrows) (close-ups from **(a)**, areas in squares). Scale bars: **a,b,d** 20 μm; **c,e**, 10 μm.

## Discussion

FeSODs were long believed to be chloroplast proteins involved in superoxide scavenging during photosynthesis. However, the scavenging capacity of *Arabidopsis* FSD1 was challenged, because its transcript levels remained unchanged in response to many environmental conditions^9,12,25,26^. Here, we show for the first time that FSD1 is localized not only in plastids, but simultaneously also in the nuclei and cytoplasm of *Arabidopsis* cells. Moreover, FSD1 relocalizes to the plasma membrane under salt stress conditions.

### FSD1 might protect root proliferation activity under adverse environmental conditions

Using translational fusion constructs with native promoter, GFP-tagged FSD1 exhibited a tissue-specific expression pattern in *Arabidopsis* root tip. This indicates that FSD1 may also have developmental roles that are conditionally determined. Hence, FSD1 might be involved in the regulation of the redox status in dividing cells, like root initials. It is known that the root meristematic activity as well as the quiescent center organization is maintained by redox homeostasis which acts downstream of the auxin transport ^26-29^. Intriguingly, FSD1 tissue-dependent expression pattern largely correlates with auxin maxima in the root tip^30,31^, as well as with superoxide anion maxima^32^. Furthermore, endodermis formation requires *SCARECROW (SCR)* and *SHORTROOT (SHR)*, two GRAS-type transcription factors, expressed in the endodermis/cortex initials and quiescent center^33,34^. FSD1 might also contribute to the regulation of SCR and SHR, which is supported by the high expression of *FSD1* in fluorescence-activated cell sorting (FACS)-isolated protoplasts expressing endoplasmic reticulum targeted GFP under the control of the SCARECROW promoter^35^. This expression was elevated in salt-stressed protoplasts. Considering our results about the role of FSD1 in salt stress tolerance, FSD1 may be involved in the maintenance of redox homeostasis in the endodermis/cortex initials of the root tip.

### FSD1 is required for *Arabidopsis* response to the salt stress

Our localization data suggest that FSD1 functions are not only restricted to the cytoplasm and plastids, because we provide here the first evidence on the nuclear localization of superoxide dismutase in plants. It was previously found that mammalian SOD1 is rapidly relocated to the nucleus upon H_2_O_2_ triggered oxidative stress36. In this case, SOD1 binds to specific DNA nucleotide sequences and triggers the expression of genes involved in oxidative resistance and DNA repair. It may also bind to and regulate the stability of specific mRNAs^36^. SOD1 nuclear functions are unrelated to its catalyzing of superoxide removal^37^. Nucleotide sequences of *FSD1* as well as structure of FSD1 catalytic and other domains differ considerably from SOD1^10^. Thus, the nuclear function of FSD1 cannot be easily anticipated, but it certainly deserves further study.

The localization of FSD1 to chloroplasts is determined by an N-terminal transit peptide identified previously^13^. According to comparative studies of three *Arabidopsis* isoforms, FSD1 is crucial neither for chloroplast integrity^12^, nor for cell protection under photooxidative stress^26^. It is likely that the protective role of FSD1 depends on the severity of the external conditions and might be triggered under harsh stress conditions. The protective roles of FSD1 were reported in transgenic tobacco and maize, where overexpression of this enzyme in chloroplasts enhanced the efficiency of thylakoid and plasma membrane protection^14,15^. Our results suggest that FSD1 is important for *Arabidopsis* germination under salt stress and salt stress tolerance. As indicated by the salt stress response of the complemented lines, cytosolic and likely also nuclear FSD1 pools are crucial for the acquisition of full tolerance to salinity during germination. Altogether, our results emphasize the importance of FSD1 in the regulation of cytosolic and also possibly nuclear redox homeostasis in response to salinity stress.

### Salt-induced relocation of FSD1 to the plasma membrane and periplasmic ROS production

Plasmolysis is a primary consequence of salt (osmotic) shock in plants^38,39^. Our data showed strong accumulation of FSD1 in Hechtian strands and reticulum during plasmolysis. These tubular structures are plasma membrane extensions providing a physical connection between the retracted protoplasts and the cell wall^38^. The protoplast shrinkage and formation of Hechtian strands is accompanied by rapid plasma membrane remodeling and modifications^40^, likely driven by the documented generation of ROS, which are known to affect the plasma membrane properties by lipid peroxidation^41^. Therefore, we suggest that FSD1 is relocated to the plasma membrane during salt shock in order to control ROS-mediated plasma membrane modifications in Hechtian strands and reticulum during plasmolysis. Such function might be assigned also to thioredoxin H9 which has similar periplasmic localization^42^. The plasma membrane localization of FSD1, which was also experimentally confirmed in several proteomic studies^43-45^, is most likely mediated by two predicted hydrophobic helices^46^. The EEFNAAAATQFGAGWAWLAY region was significantly predicted to have an extracellular orientation (Supplementary Fig. 14).

### FSD1 is likely involved in endosperm rupture during seed germination

Seed germination is a complex process encompassing multiple events governed by tight phytohormonal regulation. Micropylar endosperm represents the last mechanical barrier constraining the radicle emergence. Endosperm rupture is preceded by its weakening, controlled by the inhibitory effect of abscisic acid (ABA) and promoting effect of ethylene^47^. Furthermore, ROS contribute to this process by oxidizing the cell wall polysaccharides and subsequent cell wall loosening^48^. Here, we provide data showing FSD1 upregulation and local accumulation in the micropylar endosperm during endosperm weakening and rupture, which is subsequently decreased after primary root emergence. Such accumulation of FSD1-GFP at the micropylar endosperm before and during endosperm rupture by emerging radicle indicates that it may be involved in the local catalysis of superoxide conversion to hydrogen peroxide. Indeed, *FSD1* shows unique transcriptional changes during seed germination in comparison to other SOD isoforms^48^, supporting the specific role of FSD1 during endosperm weakening and rupture.

In summary, we show developmentally regulated tissue-specific expression pattern, triple subcellular localization and provide evidence for the new role of FSD1 in the salt stress, which is unique among plant SODs. These new features make FSD1 favorable candidate for potential biotechnological applications.

## Author Contributions

PD, YK, JB, VZ and MO performed the experiments and analyses. TT coordinated the experiments, supervised the project and helped with data assessment. JŠ provided the infrastructure and helped with the interpretation of the results. PD, YK and TT drafted the manuscript which was revised and edited by MO and JŠ. All authors approved the final version of the manuscript.

## Acknowledgements

This research was funded by Grant No. 19-00598S from the Czech Science Foundation GAČR and by the ERDF project “Plants as a tool for sustainable global development” (No.CZ.02.1.01/0.0/0.0/16_019/0000827). We thank to Dr. O. Šamajová for her valuable advices during microscopic observations. We are also thankful to Michael Wrzaczek for critical reading of the manuscript.

## Data availability

Data from this study are available within the paper and the Supplementary Information or from the corresponding authors upon request.

