## Supplementary Table and Figures for "FSD1: a plastidial, nuclear and cytoplasmic enzyme relocalizing to the plasma membrane under salinity"

| Oligo name | Sequence 5' - 3' | Use |
|---|---|---|
| C-terminal fusion: Promoter + gene_FWD | GGGGACAACTTTGTATAGAAAAGTTGCG GGCATTTGCTGTATATGAGAAGCC | PCR |
| C-terminal fusion: Promoter + gene_REV | GGGGACTGCTTTTTTGTACAAACTTGG AGCAGAAGCAGCCTTGGC | PCR |
| C-terminal fusion: 3´UTR_FWD | GGGGACAGCTTTCTTGTACAAAGTGGTC GCAAATTTCTGAACAATTTGAC | PCR |
| C-terminal fusion: 3´UTR_REV | GGGGACAACTTTGTATAATAAAGTTGG TTACTTCAAAGCTCACTCTCTGC | PCR |
| N-terminal fusion: Promoter_FWD | GGGGACAACTTTGTATAGAAAAGTTGGC GCTGTATATGAGAAGCCCTTC | PCR |
| N-terminal fusion: Promoter_REV | GGGGACTGCTTTTTTGTACAAACTTGG TCTTTGTAATTGAAGCTGCAC | PCR |
| N-terminal fusion: Gene + 3´UTR_FWD | GGGGACAGCTTTCTTGTACAAAGTGGGA ATGGCTGCTTCAAGTGCTG | PCR |
| N-terminal fusion: Gene + 3´UTR_REV | GGGGACAACTTTGTATAATAAAGTTGC CTCACTCTCTGCATTGGTTCGTC | PCR |
| qFSD1_FWD | TCACCGCAAACTACGTCCTC | qPCR |
| qFSD1_REV | TCATATGCGGCTCCAAAGCA | qPCR |
| qEF1α_FWD | TGAGCACGCTCTTCTTGCTTTCA | qPCR |
| qEF1α_REV | GGTGGTGGCATCCATCTTGTTACA | qPCR |

**Supplementary table 1. List of used primers.**

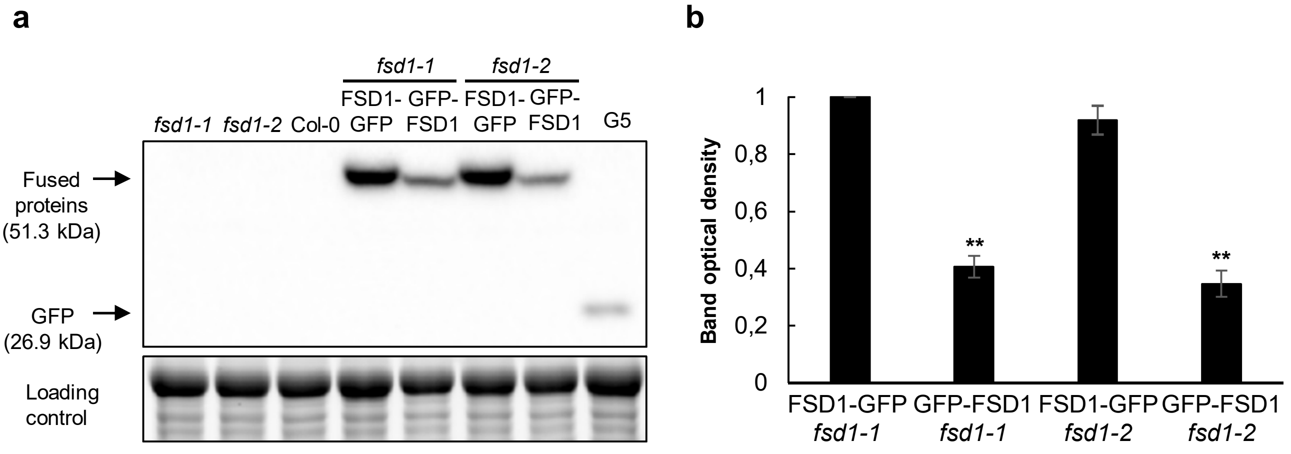


**Supplementary Fig. 1. Detection of GFP fused FSD1 in complemented *fsd1* mutants expressing *FSD1-GFP* and *GFP-FSD1* using anti-GFP antibody. a**, Immunoblots showing the presence of GFP fused FSD1 in 14-days-old complemented *fsd1* mutants expressing *proFSD1::FSD1:GFP* (FSD1-GFP) or *proFSD1::GFP:FSD1* (GFP-FSD1), and the absence of signal in Col-0 and *fsd1* mutants. G5 line expressing 35S::GFP were used as positive control. **b**, Quantification of optical density of band in **(a)**. The densities are expressed as relative to the highest value. Error bars represent standard deviation. Stars indicate statistically significant difference between GFP-FSD1 and FSD1-GFP abundance (one-way ANOVA, $^{**}p < 0.01$).

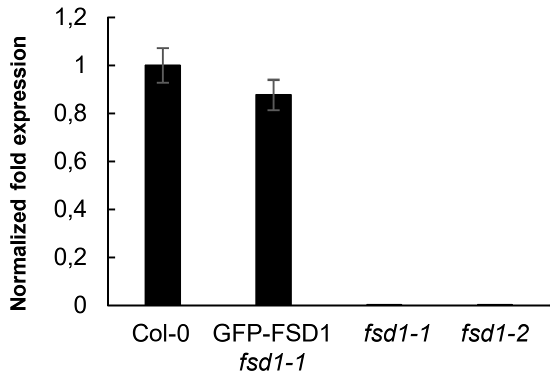


**Supplementary Fig. 2. Quantification of *FSD1* transcript level in 14-day-old Col-0, *fsd1* mutant seedlings and seedlings of complemented *fds1-1* mutant expressing *proFSD1::GFP:FSD1* (GFP-FSD1).**

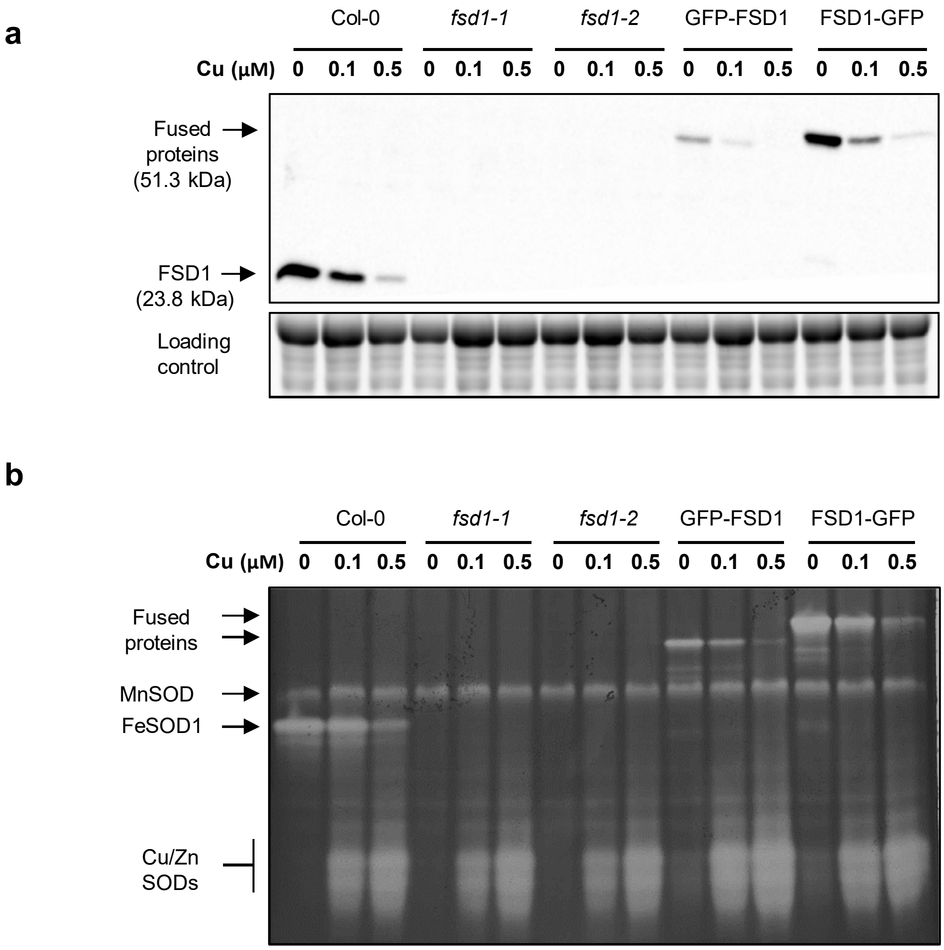


**Supplementary Fig. 3. SOD isoform abundances and activities in *fsd1* mutants and complemented *fsd1* mutant expressing *proFSD1::FSD1:GFP* (FSD1-GFP) or *proFSD1::GFP:FSD1* (GFP-FSD1) in response to different Cu concentrations a,** Immunoblot probed with anti-FSD1 antibody. **b**, visualization of activities of SOD isoforms on native polyacrylamide gels.

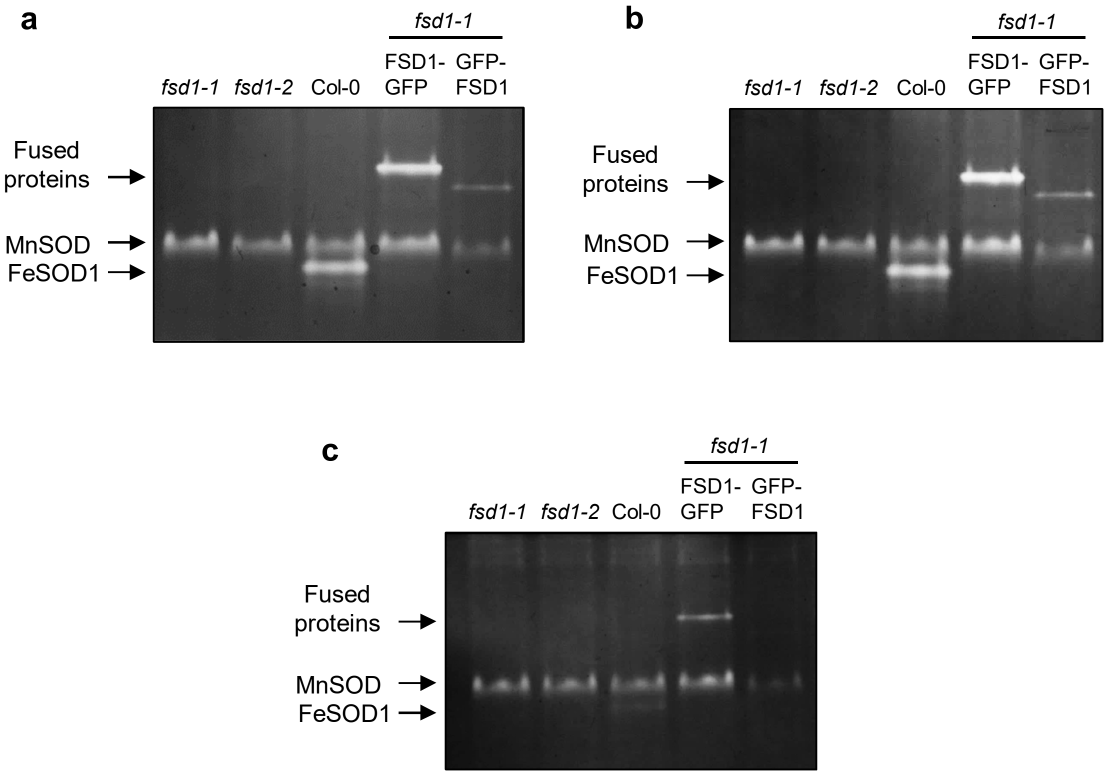


**Supplementary Fig. 4. Identification of SOD isozymes in complemented *fsd1* mutant expressing *proFSD1::FSD1:GFP* (FSD1-GFP) or *proFSD1::GFP:FSD1* (GFP-FSD1) by specific inhibitors**. **a-c,** Visualisation of SOD isozymes on native polyacrylamide gel whithout preincubation **(a)**, with preincubation in KCN inhibiting Cu/ZnSODs activity **(b)** and with preincubation in $H_2O_2$, inhibiting FeSOD and Cu/ZnSODs **(c)**.

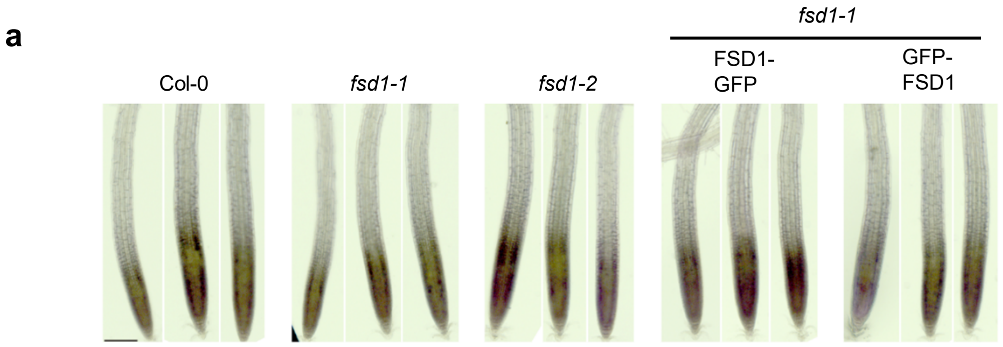


**Supplementary Fig.5. Histochemical detection of superoxide. a**, Specific staining of superoxide on roots of 7-days-old Col-0, *fsd1-1*, *fsd1-2* mutants and *fsd1-1* mutants *proFSD1::FSD1:GFP* (FSD1-GFP) or *proFSD1::GFP:FSD1* (GFP-FSD1) by using nitro blue tetrazolium chloride.

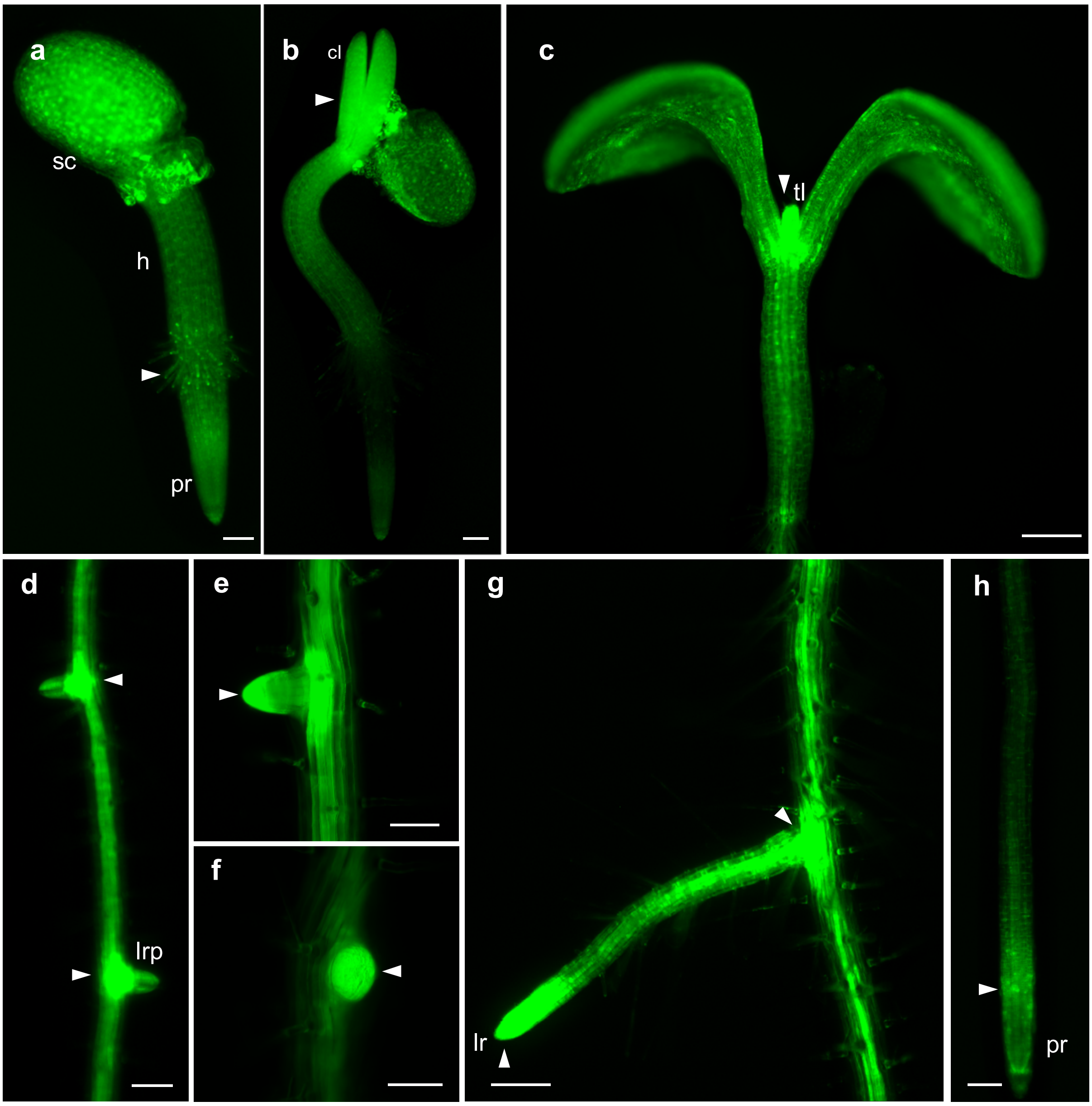


**Supplementary Fig. 6. Localization of FSD1-GFP visualized by ZOOM stereomicroscope at different developmental stages of young *Arabidopsis* seedlings**. **a**, just emerged seedling at 1st day after germination (DAG) grown out of a seed coat (sc) with hypocotyl (h) and primary root (pr). **b**, cotyledons (cl) released from a seed coat at 2nd DAG. **(c)**, fully opened cotyledons and emerging first true leaves (tl) at 5th DAG. **d-f**, lateral root primordia (lrp), 6th DAG. **g**, fully developed elongating lateral root on 8th DAG. **h**, primary root on 8th DAG. Arrowheads point to the highest intensity of GFP signal. Scale bars: **a**, **b**, **d**, **e**, **f**, **h**, 50 µm**;** **c**, **g**, 200 µm.

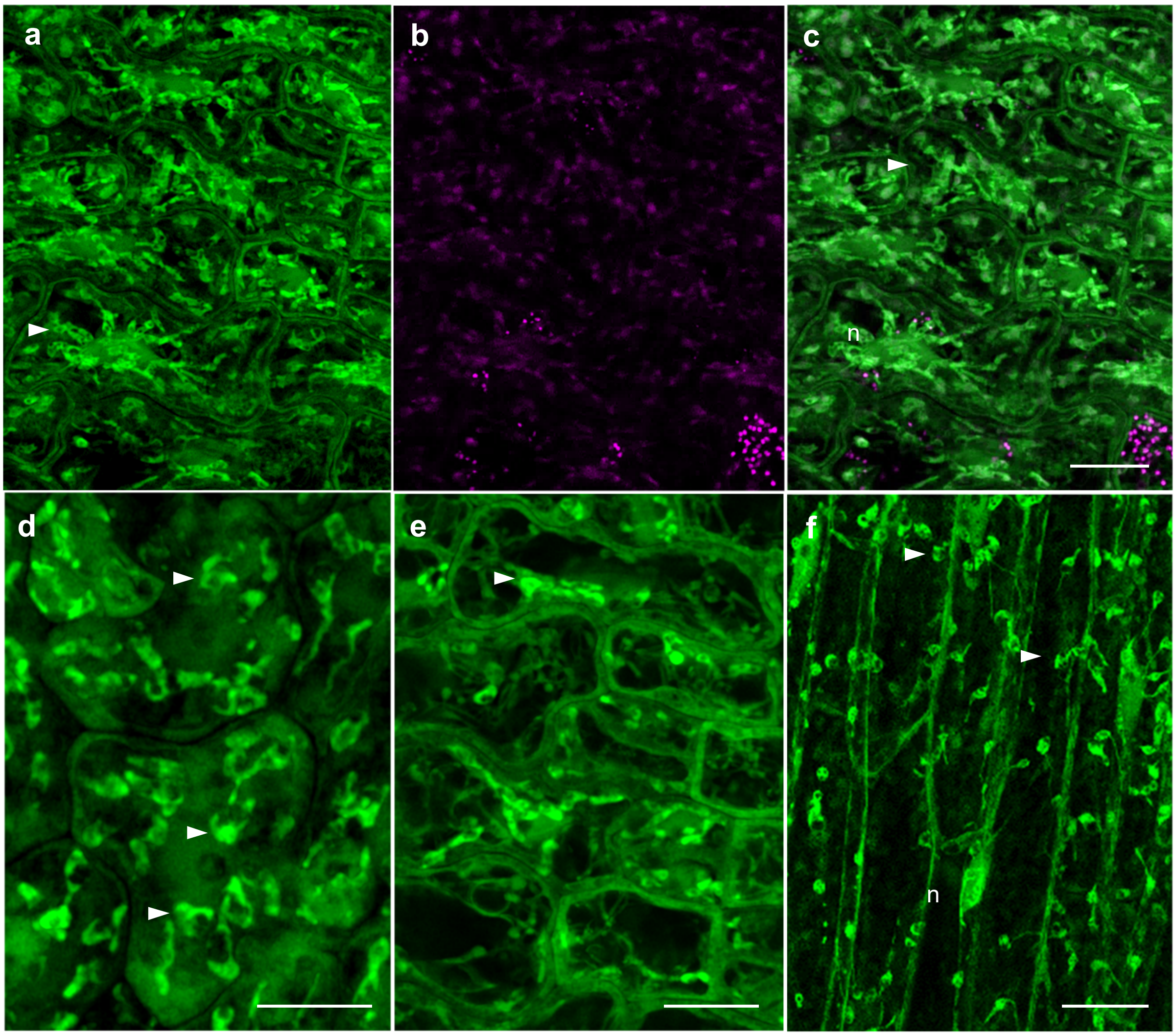


**Supplementary Fig. 7. FSD1-GFP localization in epidermal cells of etiolated *Arabidopsis* seedlings observed using Airyscan confocal laser scanning microscope. a-e**, cotyledons on 2nd day after germination (DAG)**. f**, hypocotyl on 2nd DAG. Arrowheads indicate plastids, (n) stands for nuclei. Channels: green - FSD1-GFP; magenta - chlorophyll *a* autofluorescence. Scale bars: **a-e**, 10 µm; **f**, 20 µm.

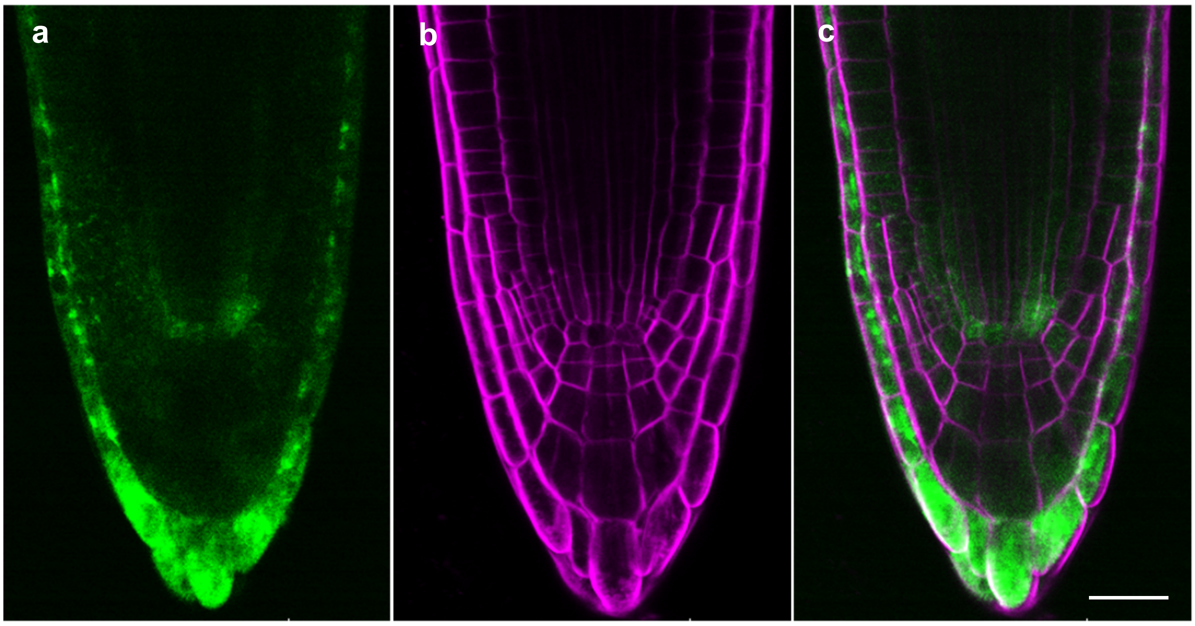


**Supplementary Fig. 8. Overview of FSD1-GFP tissue-specific localization in primary root apex with root cap, QC and part of the meristem**. **a-c**, Seedlings at 1st day after germination expressing FSD1-GFP were stained with FM4-64 and observed under confocal laser scanning microscope. Channels: green - FSD1-GFP **(a)**; magenta - FM4-64 **(b)**; overlay **(c)**. Scale bar: 20 µm.

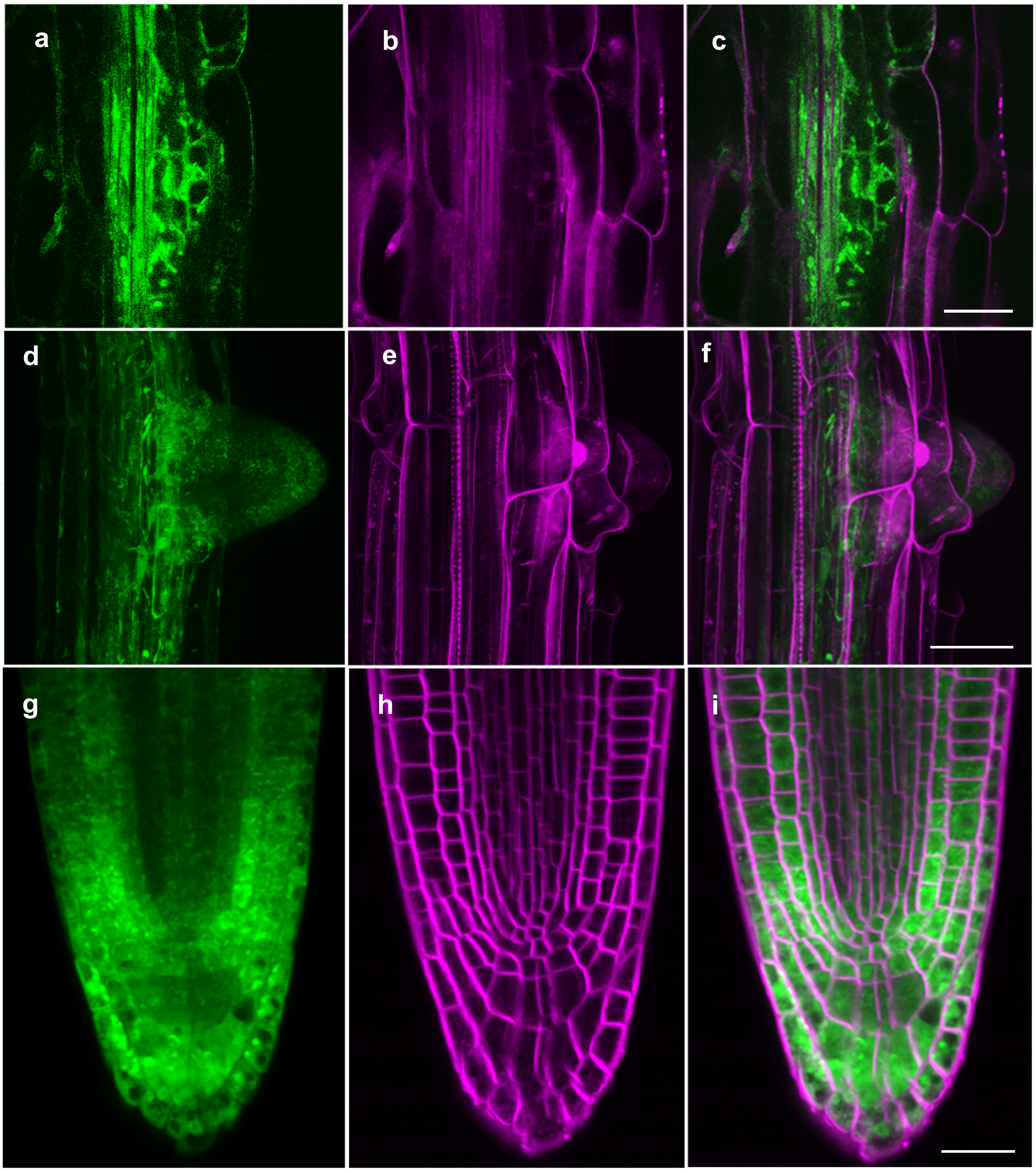


**Supplementary Fig. 9. Tissue-specific expression of FSD1-GFP during *Arabidopsis* lateral root development. a-i**, Lateral root primordia and lateral roots of seedlings at 6th and 8th day after germination (DAG) were observed by confocal laser scanning microscopy. Lateral root primordia emerging from the pericycle cells **(a-c)** at meristem formation and quiescence center specification stage (VI) (6th DAG). Lateral root primordia at meristem formation stage **(d-f)**. Lateral root tip **(g-i)** with root cap and meristem (8th DAG). Channels: green - FSD1-GFP; magenta - FM 4-64. Scale bars: 20 μm.

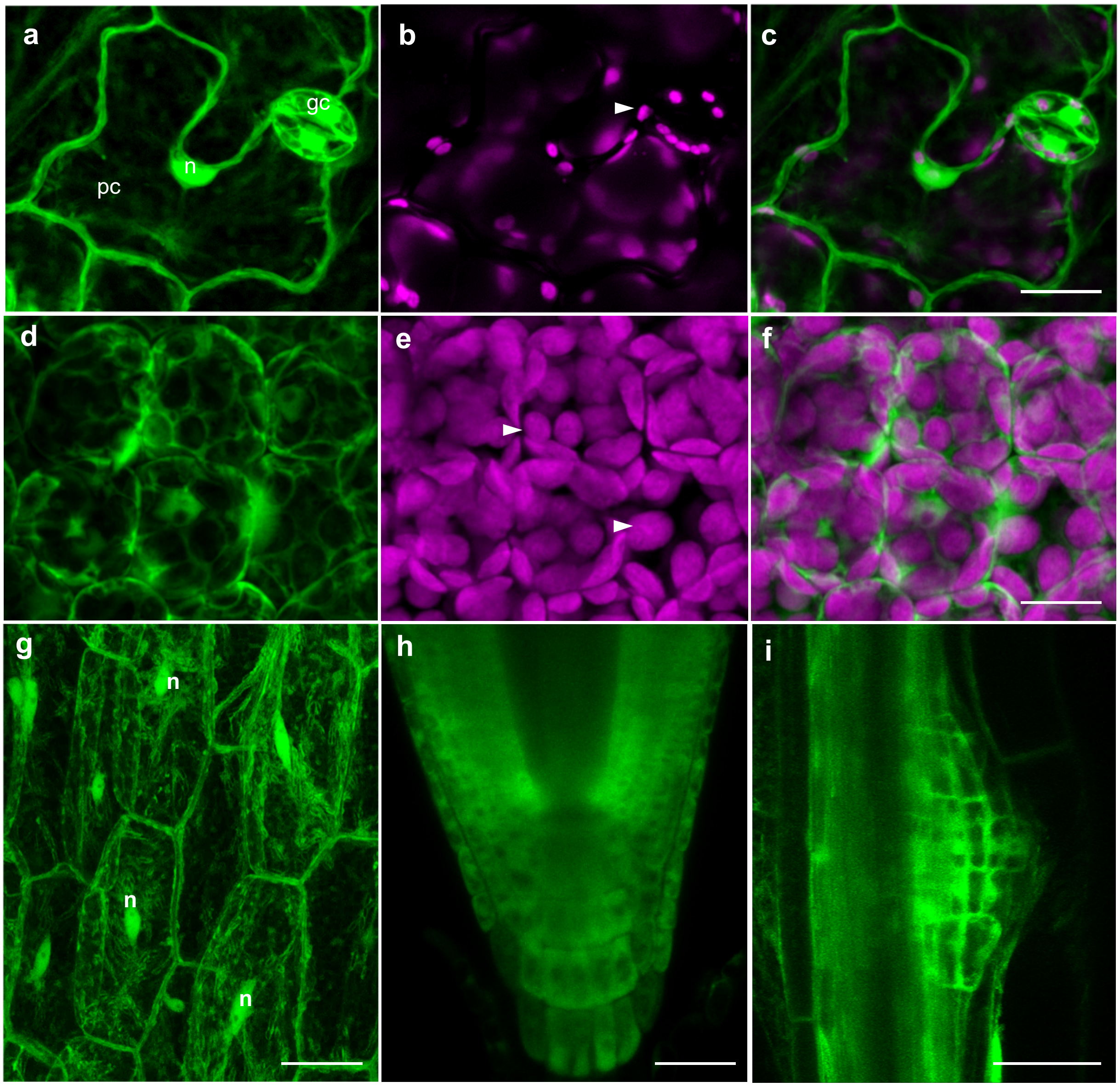


**Supplementary Fig. 10. Overview of GFP-FSD1 localization in cells of above- and underground organs of *Arabidopsis* seedlings revealed by Airyscan confocal laser scanning microscopy**. **a-c**, adaxial surface of cotyledon with pavement (pc) and guard (gc) cells. **d-f**, mesophyll cells. **g**, hypocotyl cells. **h**, primary root apex. **i**, emerging lateral root. Arrowheads indicate plastids. (n) stands for nuclei. Channels: green - FSD1-GFP; magenta – chlorophyll *a* autofluorescence. Scale bars: **a-f**, 10 µm; **g-i** 20 µm; .

**a**

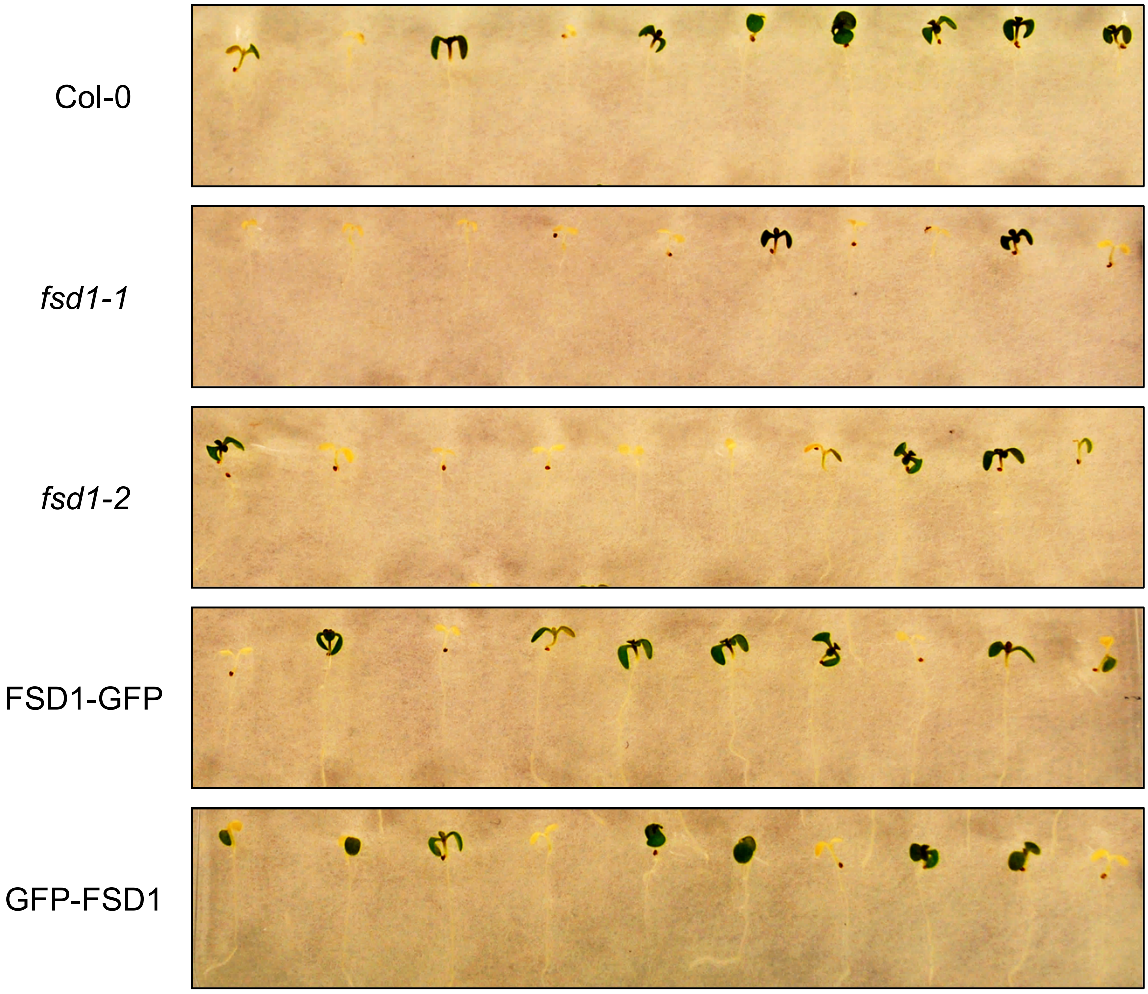


**Supplementary Fig. 11. Salt stress response of Arabidopsis lines with modified *FSD1* expression.** **a**, Representative pictures of 9-day-old Col0, *fsd1* mutants and *fsd1-1* mutants expressing *proFSD1::FSD1:GFP* (FSD1-GFP) or *proFSD1::GFP:FSD1* (GFP-FSD1) seedlings exposed to 150 mM NaCl for 5 days.

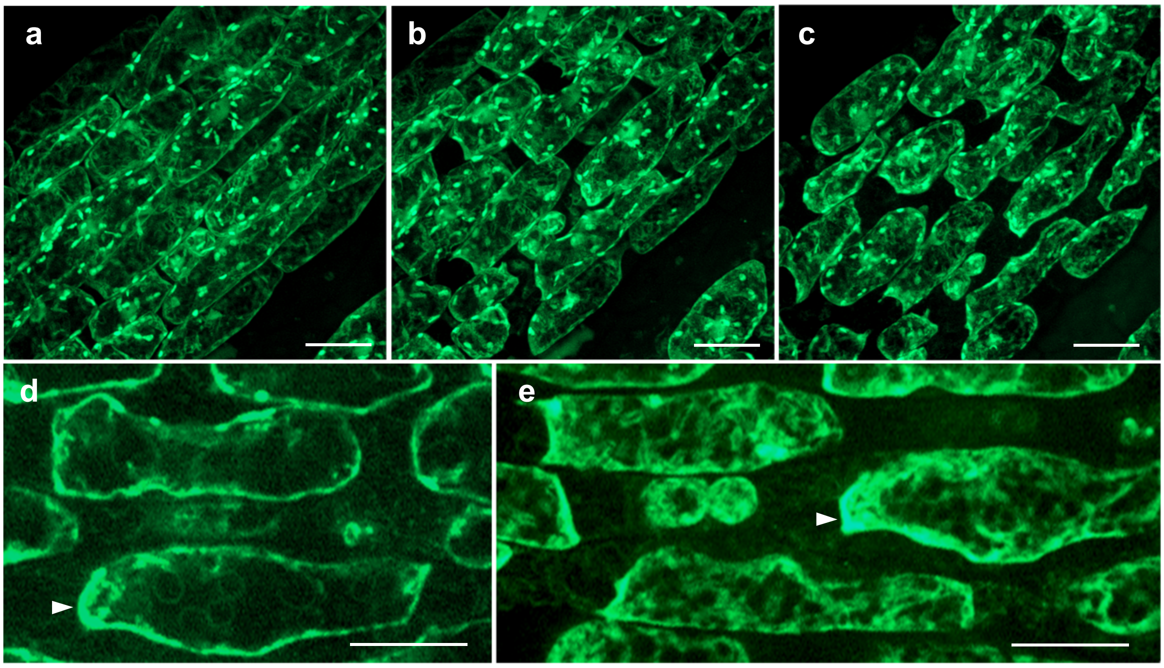


**Supplementary Fig. 12. Plasmolysis of hypocotyl epidermal cells induced by salt stress.** The progression of both concave and convex plasmolysis induced by 500 mM NaCl was observed using spinning disc microscope on *fsd1-1* mutants expressing *proFSD1::FSD1:GFP* (FSD1-GFP). **a**, intact cells; **b**, 5 min treatment; **c**, 30 min treatment; **d-e**, magnification of plasmolyzed cells. Arrowheads point to FSD1-GFP signal accumulation. Scale bar: 20 μm.

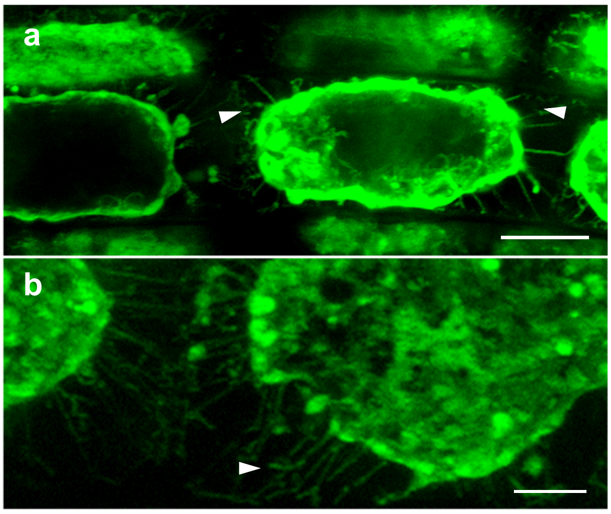


**Supplementary Fig. 13. Salt stress response in *Arabidopsis* primary roots.** Plasmolysis was induced by the treatment of seedlings with liquid ½ MS medium containing NaCl (250 mM) for 15 min. **a,** concave plasmolysis and Hechtian strands (arrowheads) in epidermal cells of primary root elongation zone of complemented *fsd1-1* mutant harboring FSD1-GFP. b, detail of Hechtian strands. Scale bars: 10 µm.

### TMpred

Home | Contact

### TMpred output for UNKNOWN

**[EMBnet-Server]** Date: Sun Feb 23 15:34:21 2020

---

```
TMpred prediction output for : TMPRED.25035.3302.seq

Sequence: MAA...ASA   length:     212
Prediction parameters: TM-helix length between 17 and 33


1.) Possible transmembrane helices
==================================
The sequence positions in brackets denominate the core region.
Only scores above  500 are considered significant.

Inside to outside helices :   2 found
      from        to    score center
 124 ( 124) 141 ( 141)    815    133
 151 ( 151) 171 ( 171)    531    161

Outside to inside helices :   2 found
      from        to    score center
 124 ( 124) 141 ( 141)    489    133
 153 ( 153) 171 ( 171)    538    161



2.) Table of correspondences
============================
Here is shown, which of the inside->outside helices correspond
to which of the outside->inside helices.
  Helices shown in brackets are considered insignificant.
  A "+"  symbol indicates a preference of this orientation.
  A "++" symbol indicates a strong preference of this orientation.

           inside->outside | outside->inside
   124- 141 (18)  815 ++   |(  124- 141 (18)  489    )
   151- 171 (21)  531      |   153- 171 (19)  538


3.) Suggested models for transmembrane topology
===============================================
These suggestions are purely speculative and should be used with
EXTREME CAUTION since they are based on the assumption that
all transmembrane helices have been found.
In most cases, the Correspondence Table shown above or the
prediction plot that is also created should be used for the
topology assignment of unknown proteins.

2 possible models considered, only significant TM-segments used

*** the models differ in the number of TM-helices ! ***

-----> STRONGLY prefered model: N-terminus inside
 2 strong transmembrane helices, total score : 1353
 # from   to length score orientation
 1  124  141 (18)     815 i-o
 2  153  171 (19)     538 o-i

------> alternative model
 1 strong transmembrane helices, total score : 538
 # from   to length score orientation
 1  153  171 (19)     538 o-i
```

**Supplementary Fig. 14. Prediction of transmembrane domains in aminoacid sequence of FSD1 according to Tmpred server (Hofmann and Stoffel, 1993).**
